# Structural–functional calibration corrects single-neuron identity errors in volumetric calcium imaging

**DOI:** 10.64898/2026.08.20.745679

**Authors:** Xiang Liu, Dongzhou Gou, Chao Song, Junjie Zhao, Mianzhi Liu, Siyuan Rao, Ye Liang, Linlu Xu, Heng Mao, Yanmei Liu, Junwen Wang, Lei Ma, Haoyu Li, Changliang Guo, Liangyi Chen

## Abstract

Volumetric calcium imaging is increasingly used to capture larger neuronal populations at higher throughput, but high-speed axial sampling can compromise single-neuron identity. Here we identify **cross-plane identity duplication** as a structured error in volumetric imaging: anisotropic axial blurring and plane-wise functional segmentation can repeatedly detect the same neuron across adjacent planes, creating **duplicate functional nodes** that inflate neuronal counts and **distort network phenotypes**. We developed **Comprehensive Label-Guided (CLG) volumetric imaging**, a structural–functional calibration framework that **uses nuclear labels as stable three-dimensional identity anchors for calcium signals**. CLG combines nuclear labeling, deep-learning-based 3D segmentation, anatomical registration and identity-guided trace reassignment. In larval zebrafish whole-brain recordings, CLG resolved ∼30,000 redundant detections and reduced estimated neuronal counts by 37–46%. In mouse visual cortex, CLG consolidated ∼40% of putative duplicates and recovered over 2,000 active neurons missed by calcium-only analysis. Across baseline and perturbed conditions, calibration stabilized graph-derived measurements of hub organization, long-range correlations and network resilience. CLG therefore defines **an anatomy-constrained identity-calibration layer** for reliable single-neuron-resolved volumetric imaging.

## Main

Volumetric calcium imaging is increasingly used to measure neuronal activity across large brain volumes, enabling circuit-level analysis in systems ranging from larval zebrafish to mammalian cortex^1–4^. However, the reliability of such analyses depends on a basic assumption: each detected functional region corresponds to a unique neuron. In high-throughput volumetric imaging, this assumption can fail. Rapid acquisition is commonly achieved by sparse axial sampling or by optical designs that extend the effective sampling volume^5–11^, both of which increase the impact of the anisotropic and depth-dependent point-spread function (PSF) ^12,13^. As volumetric platforms push toward recording more neurons, larger volumes and more axial planes per unit time, the relevant bottleneck is not only whether more neural activity can be detected, but whether each detected signal can still be assigned to the correct physical neuron. Axial blurring and depth-dependent optical degradation are well recognized in deep-tissue imaging^14–17^. As a result, fluorescence from a single soma can extend across adjacent imaging planes.

This is not random measurement noise but an identity error. When the same soma is detected across two or three axial planes, plane-wise segmentation can split one physical neuron into multiple functional units. The error inflates neuron counts, assigns the activity of one neuron to multiple functional units, and changes graph-derived measurements such as hub ranking, communicability and network robustness. These measurements are increasingly used to interpret circuit organization in disease, development and experience-dependent plasticity^18,19^. However, they depend on a basic condition: one node should correspond to one neuron.

Current calcium-imaging analysis pipelines are not designed to solve this identity problem. Calcium-only methods, including constrained non-negative matrix factorization and related algorithms^20,21^, infer neuronal footprints primarily from activity-dependent fluorescence fluctuations and therefore perform best when neuronal signals are strong and readily separable from background and neighboring sources. In dense tissue, weakly active neurons, overlapping cytosolic signals and local neuropil contamination can impair source separation, resulting in omissions, over-segmentation, or ambiguous neuronal assignment. For example, in larval zebrafish whole-brain imaging, a directly evident limitation of calcium-signal-based extraction is under-detection, with both conventional and deep learning approaches recovering fewer cells than expected in this dense imaging regime (**Supplementary Fig. 1**). Cross-plane deduplication has also been addressed heuristically; FACED2, for example, clusters ROIs across adjacent depths using spatial overlap and calcium-trace correlation ^22^. Such approaches provide useful activity-and geometry-based baselines, but their identity assignments still depend on overlap and temporal similarity rather than independent structural identity. Improved plane-wise detection therefore does not guarantee a one-to-one correspondence between detected signals and individual neurons when the same cell is sampled across multiple axial planes.

We reasoned that single-neuron identity should be defined independently of calcium activity. Neuronal nuclei provide stable three-dimensional landmarks that are less affected by instantaneous activity, neuropil fluorescence and cytosolic signal overlap. When combined with cytosolic calcium imaging, nuclear labeling can provide an anatomical scaffold for assigning functional signals to defined single-cell identities. Such a strategy should correct cross-plane duplication by merging repeated observations of the same neuron and reduce activity-dependent omission by identifying cells independently of their response amplitude.

We therefore developed Comprehensive Label-Guided volumetric imaging (CLG). The central principle is that calcium activity reports function, whereas nuclear structure defines identity. CLG uses nuclear labels as stable three-dimensional anchors, registers functional planes to this scaffold, and reassigns repeated detections to the same nucleus-defined neuron. In larval zebrafish, CLG resolved ∼30,000 redundant detections and reduced estimated neuronal counts by 37–46%. To test whether activity-and geometry-based deduplication could recover the same identities, we benchmarked a representative FACED-style ROI-chain clustering strategy against withheld CLG nucleus identities. The benchmark removed a subset of duplicate ROIs but produced substantial false splits and false merges, indicating that reducing apparent cell counts alone does not ensure correct single-neuron identity assignment. In mouse visual cortex, CLG consolidated ∼40% of putative duplicates and recovered over 2,000 active neurons missed by calcium-only analysis. These results show that structural–functional calibration converts plane-wise detections into identity-consistent single-neuron measurements.

## Results

### Nuclear labeling defines a stable three-dimensional structural scaffold in the larval zebrafish brain

We first tested whether nuclear labeling could provide a saturated structural scaffold for cell identity in the dense larval zebrafish brain. Thus we generated a dual-labeled zebrafish line, Tg(elavl3:H2B-mRuby3; elavl3:GCaMP6s) ^23^, in which neuronal nuclei were labeled with mRuby3 and cytosolic calcium signals were reported by GCaMP6s (**Fig. 1a,b; Supplementary Videos 1 and 2**). We acquired dual-channel volumetric structural images across the entire larval brain, with a nuclear channel (orange) and cytosolic GCaMP6s channel (cyan) at 1 µm axial steps (>200 Z-planes per volume, 3 s per plane, varying by individual), a sampling density that exceeded the Nyquist requirement for our axial resolution and allowed individual nuclei to be resolved across three dimensions. Rather than relying on instantaneous calcium activity to define neurons, this dataset provided an activity-independent anatomical scaffold for identifying single-cell positions throughout the brain.

**Figure 1.**
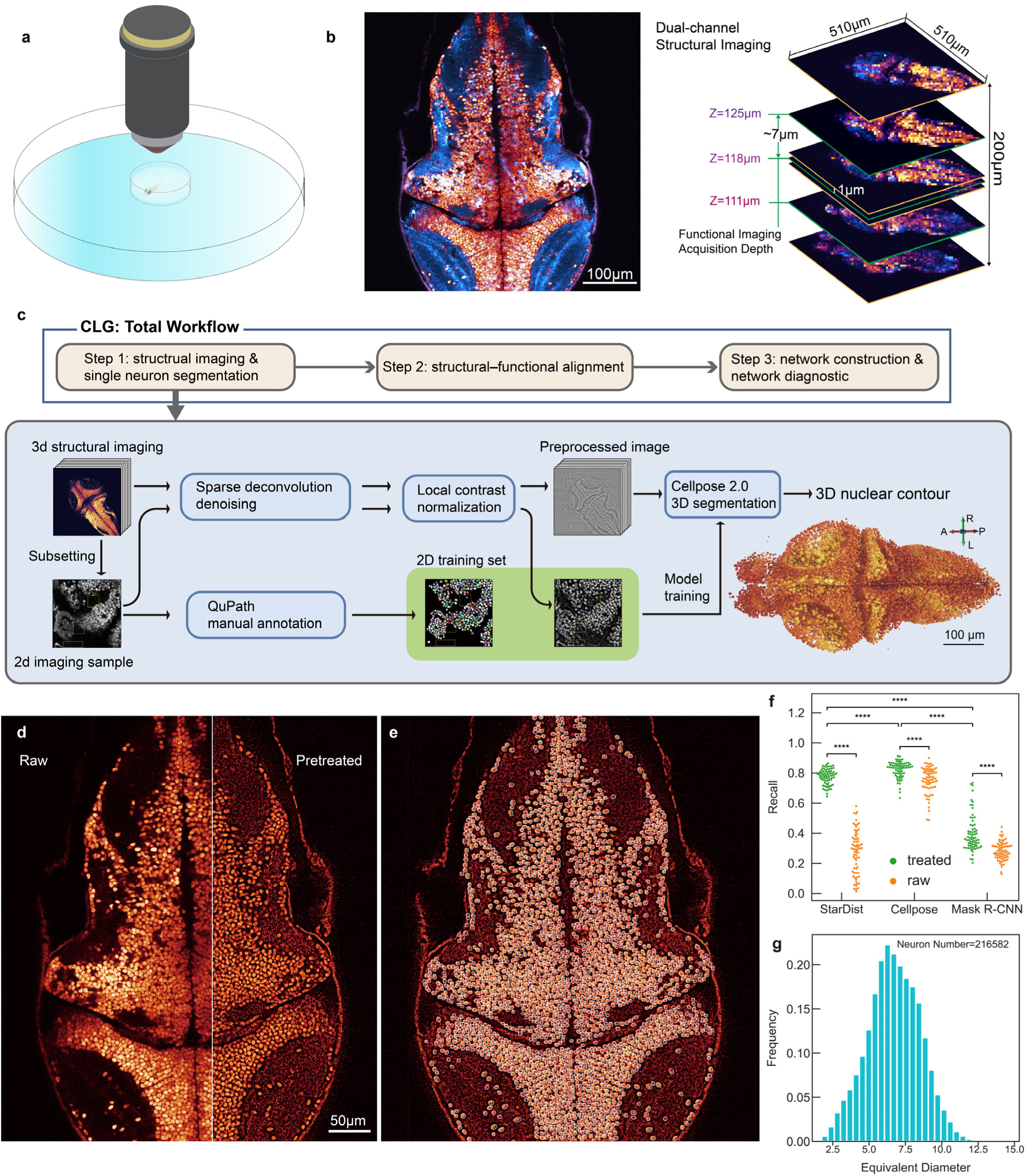
Nuclear labeling and three-dimensional segmentation define a structural scaffold for single-neuron identity in larval zebrafish. (a) Zebrafish larval imaging setup enabling stable two-photon imaging and behavioral stimulation. (b) Representative dual-channel structural images showing GCaMP6s (cyan) and mRuby3 (orange) signals (510 × 510 × 200 µm³; 1 µm Z-step; 3 s per plane), together with schematic of structural and functional axial sampling. (c) CLG workflow and 3D nuclear segmentation pipeline including denoising, local normalization, Cellpose-based segmentation, and 2D training data annotated in QuPath (scale bar, 100 µm). (d) mRuby3 nuclear images before and after preprocessing. (e) Nuclear segmentation results. (f) Effect of preprocessing on segmentation accuracy across StarDist, Cellpose 2.0, and Mask R-CNN. (g) Distribution of equivalent nuclear diameters for pooled segmented nuclei from the analyzed structural volumes.

To segment neuronal nuclei, we coupled deep learning algorithms with sparse deconvolution ^24^ and local contrast normalization (**Fig. 1c**). Sparse deconvolution improved boundary definition and signal uniformity in low signal-to-noise ratio (SNR) regions, significantly reducing fluorescence variance and enhancing the signal-to-background ratio (*P* < 0.001 for both metrics, N = 10,296 cells; **Supplementary Figs. 2–3**). Local contrast normalization (**see Methods**) further mitigated regional intensity heterogeneity caused by biological and optical factors, such as non-uniform illumination ^14^ and variable expression levels of the fluorescent transgene across different tissues ^25^, yielding uniformly high-contrast images (**Fig. 1d, Supplementary Fig. 2c,d; Supplementary Video 3**).

Following preprocessing, we evaluated three deep learning–based segmentation algorithms applied in a slice-by-slice manner: StarDist, which integrates U-Net with shape descriptors ^26^; Cellpose, a U-Net-based segmentation model that predicts spatial flow fields for instance reconstruction ^27,28^; and Mask R-CNN, a general object instance segmentation framework ^29^ (**Supplementary Fig. 4**). Ground truth datasets were generated using manual annotations created with QuPath ^30^ across 70 images spanning diverse brain regions, depths, and brightness conditions, totaling over 10,000 manually annotated nuclei (**Supplementary Fig. 5**). Models trained on preprocessed images consistently outperformed those trained on raw data (*P* < 0.0001 for all models), with Cellpose 2.0 and StarDist demonstrating high recall (mean values of 0.82 and 0.78, respectively) and significantly outperforming Mask R-CNN (mean recall: 0.40, *P* < 0.0001) (**Fig. 1e,f**). Because Cellpose 2.0 supports 3D segmentation and inter-slice association, we used it to reconstruct nuclei across volumes, enabling precise single-cell localization and contour extraction (**Fig. 1e; Supplementary Videos 4 and 5**). From these reconstructions, we estimated 54,146 ± 5,535 neurons in the zebrafish brain at 6–7 dpf (mean ± s.d., *N* = 6 larvae), with nuclei having a mean equivalent spherical diameter of 6.7 ± 1.8 µm (**Fig. 1g; see Methods**). Together, these reconstructions defined a stable three-dimensional anatomical scaffold against which functional observations could later be calibrated.

### Coarse axial sampling causes widespread cross-plane identity duplication in whole-brain zebrafish imaging

We next tested whether standard high-speed axial sampling preserves one-to-one neuron identity. To do so, we performed whole-brain functional calcium imaging of cytosolic signals using axial steps of 5–7 µm, with 7 µm used in the majority of recordings. This sampling scheme enabled rapid volumetric acquisition over ∼200 µm depth at 1 Hz, while each optical section within a 0.5 × 0.5 mm^2^ field of view was scanned in 33 ms (**Fig. 2a**). For calibration, the dual-channel structural stack provided two linked references: the mRuby3 nuclear channel defined stable three-dimensional cell identities, and the high-resolution structural GCaMP6s channel served as the registration reference for the later high-speed functional GCaMP6s recordings. Motion-corrected functional images were then registered to this structural cytosolic GCaMP6s reference, which was acquired in the same coordinate frame as the mRuby3 nuclear scaffold (**Figs. 1b, 2c; see Methods; Supplementary Video 6**). This registration chain allowed us to assign conventional plane-wise functional detections to nucleus-defined three-dimensional cell identities, enabling accurate extraction of single-cell Ca²⁺ traces (**Fig. 2b; Supplementary Fig. 6**) and direct assessment of cross-plane duplication.

**Figure 2.**
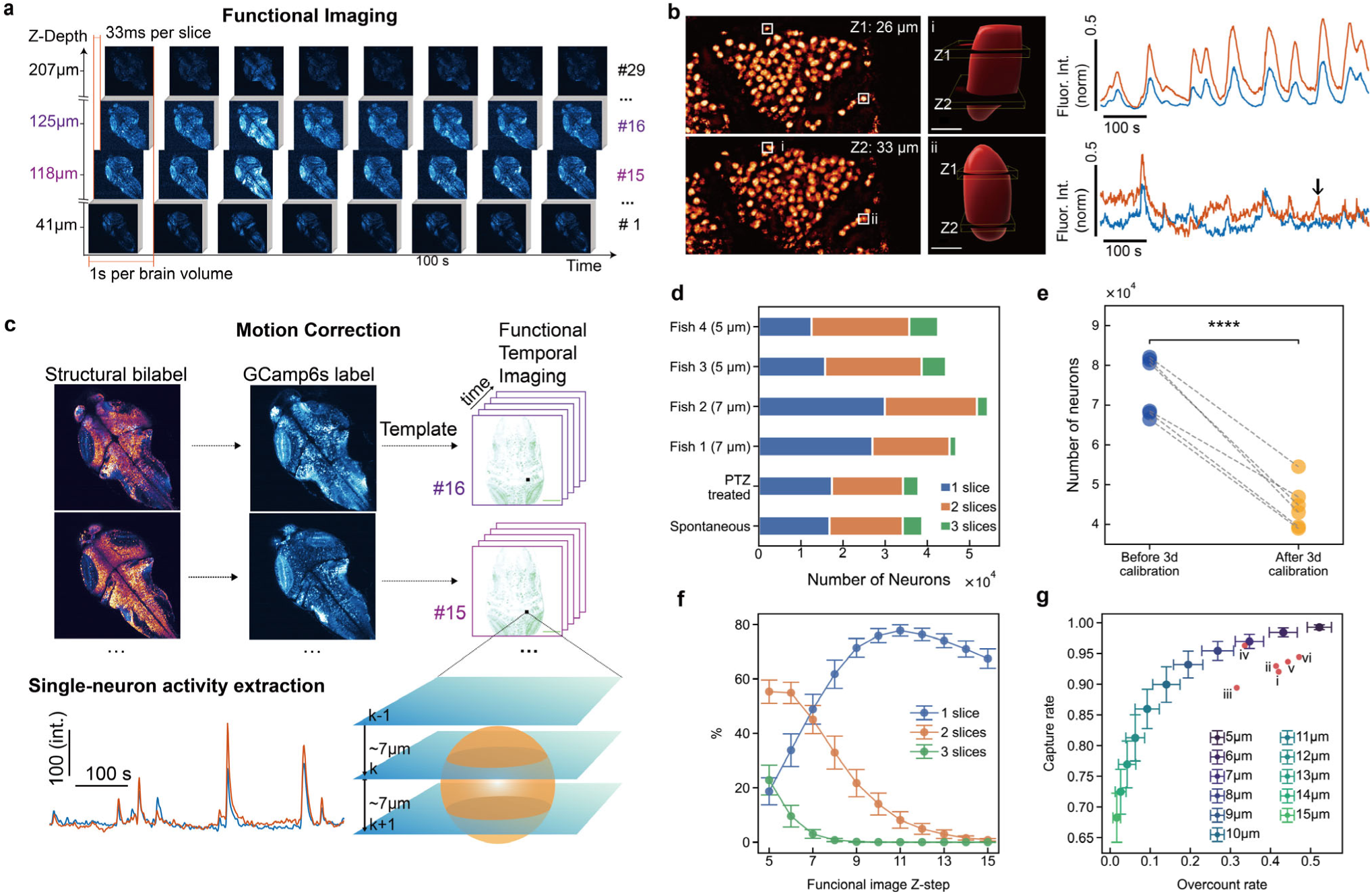
Cross-plane duplication in whole-brain functional imaging and identity-guided signal extraction. (**a**) Whole-brain GCaMP6s functional imaging (27–30 planes within 1 Hz, 200 µm depth, 5–7 µm Z-steps). (**b**) Neurons spanning multiple imaging planes. White squares (i and ii) highlight example nuclei, reconstructed in 3D (middle) to show intersection with imaging planes. Corresponding activity traces (right) reveal signal discrepancies (arrows) between planes. (**c**) Structural–functional alignment and identity-guided signal extraction. Top, alignment of functional imaging planes to the structural reference volume. Bottom, schematic example of a 3D nucleus intersecting multiple planes and the corresponding plane-specific activity traces extracted from each intersecting section. (**d**) Neurons spanning 1–3 planes across six experiments (spontaneous, PTZ; Fish 3–4 at 5 µm Z-steps, others ∼7 µm). (**e**) Reduced redundant cell counts after CLG calibration (**** ***P* < 0.0001**). (**f**) Fraction of neurons spanning multiple planes versus Z-step. (**g**) Trade-off between signal capture rate and overcount rate as a function of Z-step. Roman numerals (i–vi) mark empirical data points observed in the experiments shown in **d**: (i) Spontaneous, (ii) PTZ treated, and (iii–vi) Fish 1–4.

Despite a 7 µm step size comparable to the average neuronal diameter, many cells extended across two or more axial planes (**Fig. 2b–d; Supplementary Video 7**). Notably, Ca²⁺ signals from different segments of the same neuron could differ due to axial PSF elongation, depth-dependent SNR fluctuations, subcellular structural heterogeneity, and plane-specific neuropil contamination (**Fig. 2b, right**), indicating that functional correlation alone is insufficient for accurate cell identification ^31–33^. Across experiments, 1,000–4,000 neurons spanned three axial planes, and the number of two-plane neurons was comparable to that of single-plane neurons (**N = 6 larvae, P = 0.9542**) (**Fig. 2d; Supplementary Fig. 7**). These observations show that in rapid volumetric imaging, the problem is not simply reduced axial resolution, but loss of identity consistency across planes. Although the two-photon microscope equipped with a 1.05 NA objective provides high theoretical axial resolution ^12^, nuclei in vivo became progressively elongated along the axial dimension with imaging depth, indicating degradation of effective axial resolution in tissue (**Supplementary Fig. 8**).

Applying 2D segmentation independently to each axial plane yielded ∼80,000 putative ROIs (74,515 ± 7,528, mean ± s.d.; N = 6 larvae), consistent with prior estimates under the assumption of minimal cross-plane overlap ^10,34^. However, comparison with fully sampled structural data revealed ∼30,000 redundantly counted neurons (29,904 ± 6,112, mean ± s.d.) attributable to repeated sampling of the same cells across multiple planes (**Fig. 2e**). In practice, CLG reduced the estimated neuronal count by an average of 37.15% at 7 µm Z-steps (N = 4) and 45.74% at 5 µm Z-steps (N = 2). We then used the structural ground truth to assess sampling trade-offs: increasing axial steps beyond 12 µm reduced overcounting to less than 10% but resulted in the loss of more than 20% of detectable neurons, whereas smaller steps improved coverage but increased duplication (**Fig. 2f,g**). Thus, rapid volumetric imaging involves an inherent trade-off between overcounting and detection sensitivity. More importantly, identity duplication is a systematic consequence of high-speed volumetric sampling with coarse axial spacing, rather than a problem attributable only to a particular segmentation algorithm.

We next tested whether cross-plane duplication could be resolved by clustering with spatial proximity and temporal correlation of extracted activity ^22^. We implemented a FACED-style ROI-chain clustering benchmark on the same plane-wise ROI masks and calcium traces, while withholding CLG nucleus identities until evaluation. Using the published zebrafish FACED-style thresholds (3D distance < 20 µm and Pearson r > 0.5), the method produced 32,634 predicted clusters versus 40,375 CLG-defined nucleus identities, yielding a 19.2% undercount. More importantly, identity-level errors remained substantial: 50.2% of multi-plane CLG nuclei were split across multiple predicted clusters, 37.6% of predicted clusters merged ROIs from more than one CLG identity, and 63.9% of ROIs belonged to mixed-identity clusters (**Supplementary Fig. 9**). Across a distance–correlation threshold sweep, relaxed thresholds reduced false splits but increased false merges, whereas stringent thresholds reduced false merges at the cost of leaving true duplicates unresolved. Thus, FACED-style proximity– correlation clustering removed a subset of cross-plane duplicate ROIs but did not fully recover CLG-defined single-neuron identities. **These results show that heuristic deduplication can reduce apparent ROI counts, but it does not guarantee one-to-one single-neuron identity assignment.**

### Calibration reveals that identity duplication propagates into downstream network readouts

We next tested whether identity duplication changes downstream network readouts, rather than only neuron counts. We compared spontaneous recordings with recordings acquired 30 min after pentylenetetrazole (PTZ) treatment ^18^, using these two dynamical states to evaluate the stability of downstream readouts before and after CLG calibration (**Fig. 3a; Supplementary Fig. 10; Supplementary Videos 8–10**).

**Figure 3.**
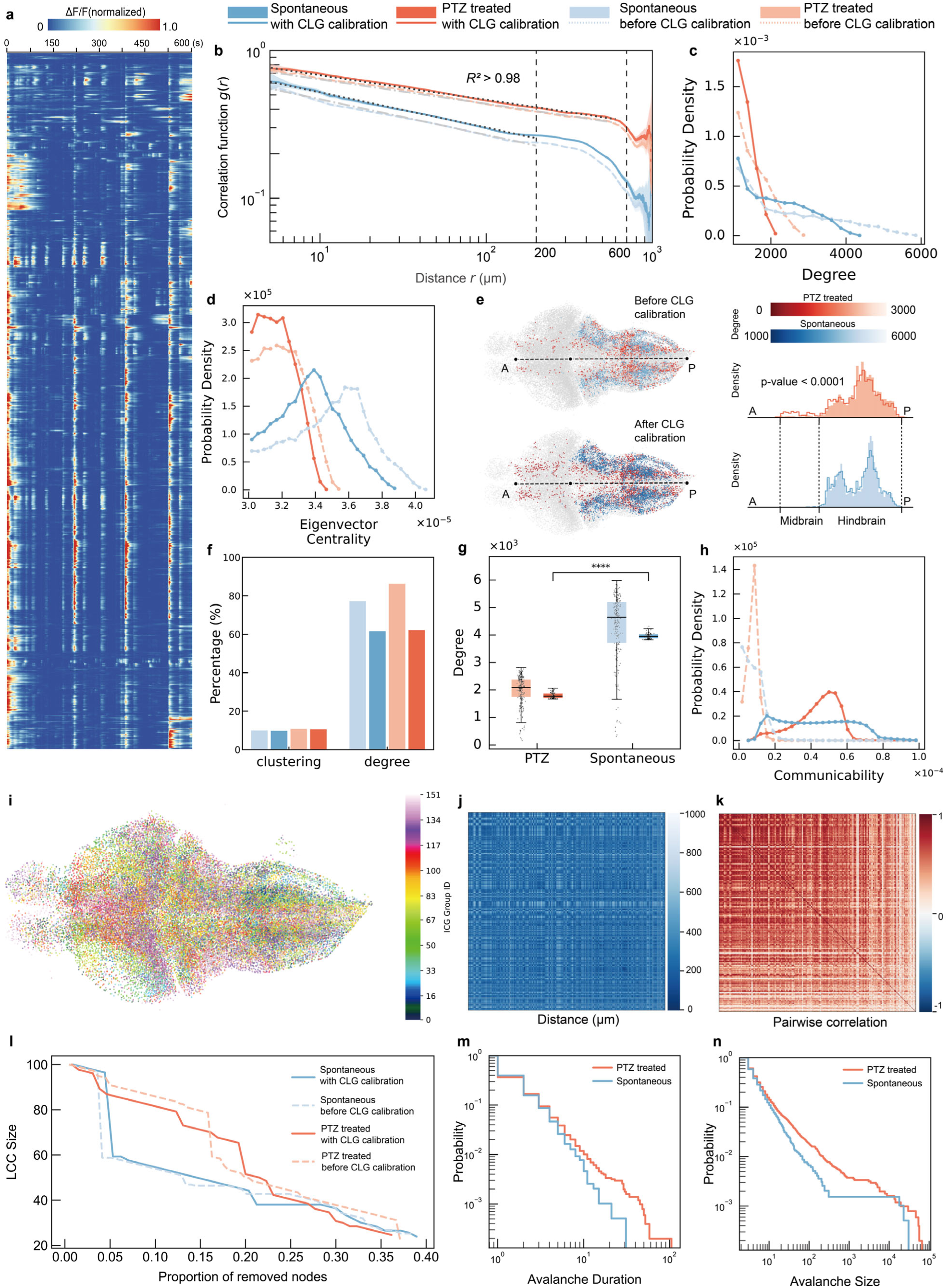
Identity duplication propagates into downstream network readouts. Network measures were compared between uncalibrated ROI-level data and CLG-calibrated neuron-identity data to assess the impact of cross-plane duplication on downstream population-level analyses. (**a**) Representative activity heatmap from PTZ-treated zebrafish datasets after denoising, principal component analysis (PCA), and ΔF/F processing. (**b**) Distance-dependent activity correlation function *g*(*r*), computed as the mean Pearson correlation between raw-activity traces of node pairs within each Euclidean-distance bin, comparing uncalibrated plane-wise ROIs and CLG-calibrated nucleus-defined identities under spontaneous and PTZ-treated conditions. (**c**,**d**) Degree (c) and eigenvector-centrality (d) distributions in uncalibrated ROI-level networks and CLG-calibrated neuron-identity networks. Dashed lines indicate uncalibrated data; solid lines indicate CLG-calibrated data. (**e)** Spatial distribution of high-degree nodes before and after calibration in PTZ-treated datasets. Marginal density plots on the right show the anterior–posterior distributions of high-degree nodes; dashed vertical lines indicate approximate midbrain and hindbrain regions, with the P value indicating the Kolmogorov– Smirnov comparison of the anterior–posterior distributions before and after calibration in the PTZ-treated dataset. (**f**) Percentage of neurons among the top 1,000 nodes ranked by degree or clustering coefficient in the CLG-calibrated networks that were also present in the uncalibrated networks. (**g**) Degree values of the top 100 highest-degree neurons after calibration compared with their corresponding values in uncalibrated datasets. Statistical analysis compares degree values between spontaneous and PTZ groups after calibration (**** ***P* < 0.0001**). (**h**) Network communicability, summarizing higher-order connectivity through multi-step paths, before and after CLG calibration (**K– S test, *P* < 0.0001**). (**i–l**) Coarse-grained network analysis and dismantling. (i) Spatial organization of coarse-grained nodes (256 cells per node) for a CLG-calibrated PTZ dataset. (j) Pairwise Euclidean distance matrix. (k) Pairwise correlation matrix ordered by coarse-grained grouping. (l) Largest connected component (LCC) size during targeted node removal using the adapted GDM algorithm. (**m,n**) Avalanche duration (m) and size (n) distributions, plotted as complementary cumulative distribution functions, for CLG-calibrated spontaneous and PTZ-treated datasets.

We first quantified distance-dependent activity correlations. We computed *g*(*r*) as the mean Pearson correlation between neuronal activity traces for cell pairs within the same Euclidean-distance bin, using uncalibrated plane-wise ROIs before calibration and nucleus-defined identities after CLG calibration (**Fig. 3b; see Methods**). PTZ treatment extended the fitted correlation range from ∼200 to ∼600 µm in both analyses, consistent with enhanced long-range coordination in seizure-like zebrafish dynamics^35,36^. However, within each condition, the uncalibrated *g*(*r*) profile was systematically lower than the calibrated profile, indicating that cross-plane duplicate ROIs altered the quantitative estimate of distance-dependent coordination.

We then used network analysis as an error-amplification test. If identity duplication were inconsequential, graph-derived measurements should remain stable after calibration. Functional networks were constructed by assigning edges between node pairs with strong activity correlations (|r| > 0.95; **see Methods**). Calibration shifted both degree and eigenvector-centrality distributions (Kolmogorov–Smirnov test, P < 0.0001 for both metrics), showing that identity duplication changed the number of high-correlation partners per node and the ranking of nodes by network influence (**Fig. 3c,d; Supplementary Fig. 11a,b**). In uncalibrated networks, degree distributions shifted rightward relative to calibrated networks, consistent with duplicate ROI nodes inflating apparent local connectivity ^4,37^.

Calibration also changed which neurons were identified as hubs. The spatial distribution of high-degree nodes differed before and after calibration (**Fig. 3e**), and the overlap of top-ranked nodes between uncalibrated and calibrated networks was incomplete, especially for clustering-coefficient rankings, a local graph metric sensitive to duplicate neighboring nodes (**Fig. 3f; Supplementary Fig. 11c**) ^38^. Similarly, the 100 highest-degree neurons after calibration showed widely variable degrees in the uncalibrated networks (**Fig. 3g; Supplementary Fig. 11d**), indicating that cross-plane duplication can alter hub assignment. Network communicability, which summarizes higher-order connectivity through multi-step paths, also changed significantly after CLG calibration (**Fig. 3h**; K–S test, P < 0.0001 for both conditions). After calibration, PTZ-treated networks showed altered hub and communicability profiles relative to spontaneous networks (**Fig. 3c,d,g,h**), with similar degree-distribution changes observed in a second biological replicate (**Supplementary Fig. 12a**). Direct visualization confirmed this reorganization (**Supplementary Figs. 12–13**). These results show that cross-plane duplication is not only a counting error. It changes the inferred nodes of the network and therefore changes graph-derived readouts of circuit organization.

To test whether identity duplication also affects graph-derived robustness estimates, we next analyzed coarse-grained functional networks. Because full single-cell networks contained more than 50,000 nodes, we grouped highly correlated neurons into super-nodes, using a coarse-graining level at which each super-node represented 256 original neurons ^39^ (**Fig. 3i–k, Supplementary Fig. 14–15; see Methods**). We then applied the adapted version of the Graph Dismantling with Machine Learning (GDM) algorithm ^40^ (**see Methods**) to remove nodes in a targeted order and measured the size of the largest connected component (LCC) during network fragmentation, with CoreHD ^41^ used as a complementary control (**Supplementary Fig. 16**). In calibrated networks, PTZ-treated datasets were more resistant to targeted node removal than spontaneous datasets: removal of the top 5% of nodes reduced the spontaneous LCC to ∼60% of its original size (**Fig. 3l and Supplementary Fig. 12b; solid blue lines**), whereas PTZ-treated networks required removal of ∼15–20% of nodes to reach a comparable collapse (**Fig. 3l and Supplementary Fig. 12b; solid red lines**). In uncalibrated networks, dismantling profiles were less consistent across animals, with PTZ networks appearing resilient in one larva but anomalously fragile in another (**Supplementary Fig. 12b; dashed lines**). Thus, cross-plane identity duplication can obscure state-dependent differences in graph-derived network robustness.

As a descriptive check on large-scale dynamics after calibration, we also examined neuronal avalanche statistics in the calibrated zebrafish datasets. In spontaneous recordings, avalanche size and duration showed broad distributions spanning multiple scales (**Fig. 3m,n**). After PTZ treatment, both distributions became heavier-tailed, with more large and long events (**Supplementary Fig. 17**), consistent with previous reports of seizure-associated large-scale dynamics ^18^. These results show that CLG-calibrated traces preserve expected state-dependent population dynamics while providing calibrated node identities for graph-based analyses.

### The same identity problem extends to mouse visual cortex

We next asked whether the same calibration logic would generalize to a markedly different physical regime. Compared with larval zebrafish whole brain, mouse visual cortex has larger neuronal somata, lower cellular density, and different scattering conditions. We imaged neuronal activity using two-photon microscopy equipped with a 25×/1.05 NA water-immersion objective (**Fig. 4a**). Neuronal nuclei from the mouse visual cortex were labeled with mRuby3 and the cytosol with GCaMP6s via adeno-associated viral delivery of one vector (**Fig. 4b**). Structural imaging with 2 µm axial steps captured the labeled nuclei within a 510 × 510 × 260 µm³ volume, identifying 7,096 labeled neurons (**Fig. 4c; Supplementary Videos 11 and 12**). Functional imaging, performed with 20 µm axial spacing—exceeding the mean neuronal diameter (∼15 µm) ^42^—yielded 10,779 putative signals prior to calibration (**Supplementary Fig. 18a)**. After CLG calibration (**Fig. 4d**), redundant detections were merged, producing 6,456 unique neurons with ∼40% consolidation and ∼10% missing rates. As in zebrafish, nuclear signals in mouse cortex were elongated along the axial dimension (**Fig. 4e**), allowing single neurons to appear in multiple functional planes (**Fig. 4d; Supplementary Video 13**). Thus, despite the markedly different anatomical and optical regime, the underlying problem remained the same: coarse axial sampling allowed single neurons to be represented multiple times across planes.

**Figure 4.**
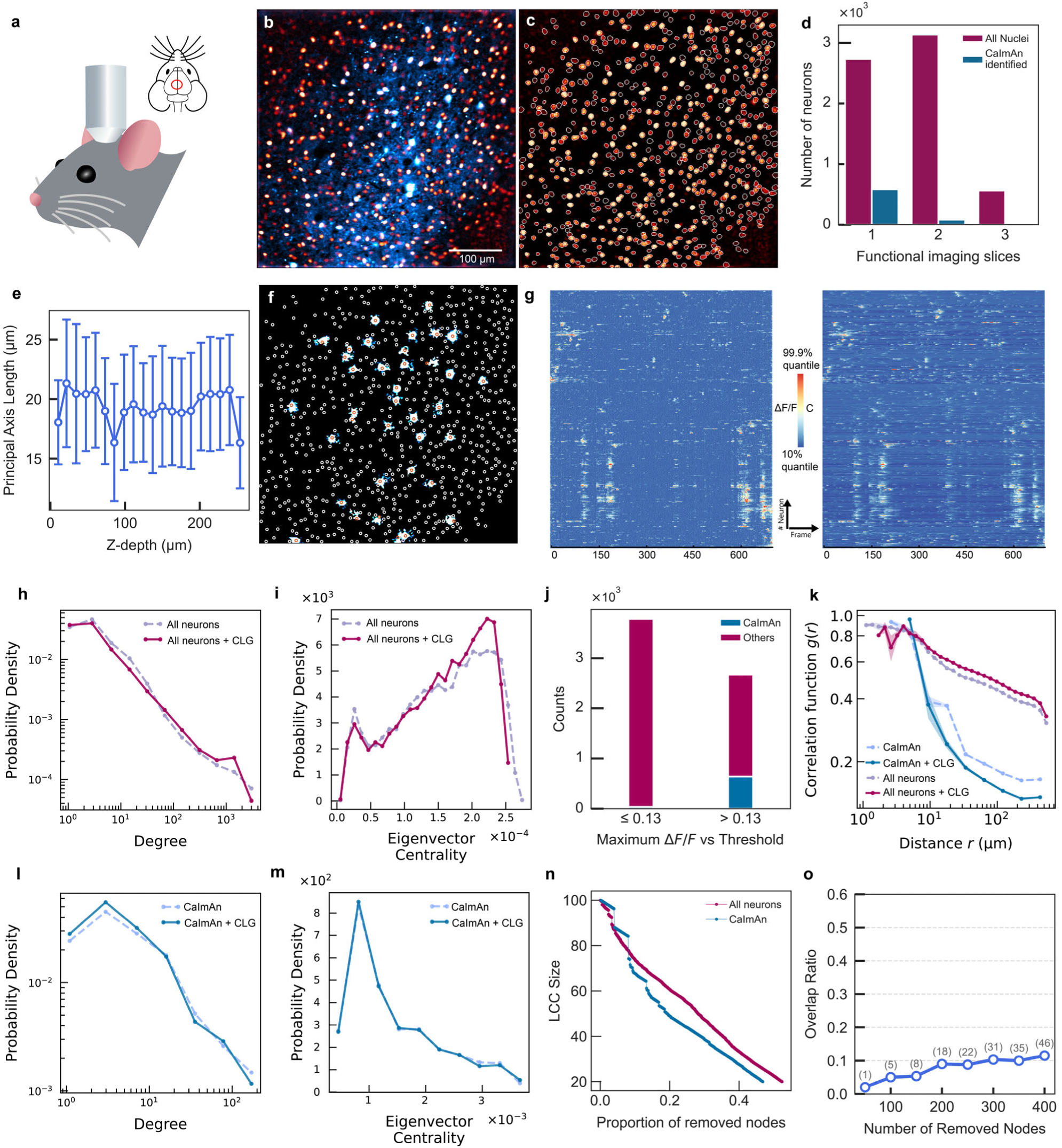
Structural-functional calibration reduces duplication and activity-dependent omission in mouse visual cortex. (**a**) Two-photon imaging setup for mouse brain imaging. (**b**) Representative dual-channel structural image (GCaMP6s, cyan; nuclear mRuby3, orange). (**c**) Example structural segmentation of neuronal nuclei in mouse visual cortex. (**d**) Number of nuclei intersecting one to three functional imaging planes; maroon, labeled nuclei identified by CLG; blue, subset also detected by CaImAn. (**e**) Principal axis length of 3D-segmented nuclei versus imaging depth (mean ± s.d., 20 Z bins). (**f**) Comparison of neuron detection by CaImAn and CLG in the same imaging plane; cyan, CaImAn components; red circles, shared detections; white circles, neurons identified only by CLG. (**g**) Comparison of Δ*F*/*F* activity traces for neurons identified by CLG (left) and corresponding CaImAn temporal components (right). (h,i) Degree (h) and eigenvector centrality (i) distributions for networks constructed from the labeled population before calibration (uncalibrated plane-wise ROIs; dashed lines) and after CLG calibration (nucleus-defined identities; solid lines). Kolmogorov–Smirnov tests confirmed significant distributional shifts (P < 0.0001 for both metrics). (**j**) Neuron classification by peak Δ*F*/*F* (threshold = 0.13), showing CaImAn-detected neurons enriched in the high-activity group. (**k**) Distance-dependent activity correlation function *g*(*r*) computed from single-cell activity, comparing calibrated and uncalibrated data for all detected neurons and CaImAn-identified neurons. (l,m) Degree (l) and eigenvector centrality (m) distributions for networks constructed from the CaImAn-identified subset before (dashed lines) and after (solid lines) CLG calibration. K–S tests showed no significant distributional shifts for either metric (**ns**). (**n**) Network dismantling using the adapted GDM algorithm, comparing LCC size for networks constructed from the full CLG-calibrated labeled population and from the CaImAn-derived subset. (**o**) Overlap of influential nodes identified during dismantling in the full CLG-calibrated network and in the CaImAn-derived network.

We then compared CLG-based segmentation with CaImAn ^20^, a widely used calcium-only analysis toolkit. CaImAn components were matched to CLG-defined nuclei using the same nearest-nucleus criterion used throughout this analysis, with detections within 30 pixels counted as matches. From motion-corrected functional data, CaImAn identified 792 putative neurons, 779 of which matched neurons detected by CLG, corresponding to 678 unique neurons after CLG calibration (**Fig. 4d,f**). Activity traces from matched neurons were highly consistent between the two methods (**Fig. 4g**; **Supplementary Fig. 18b,c**). Using a two-component Gaussian mixture model of Δ*F*/*F* distributions, we defined active neurons with a threshold of Δ*F*/*F* > 0.13 (**Supplementary Fig. 19**). Within our calibrated population, 59% of neurons were classified as silent. Among CaImAn detections, 641 neurons exceeded the activity threshold, whereas 37 fell below it. CLG recovered all 641 CaImAn-detected active neurons and additionally identified 2,032 active neurons that were not detected by CaImAn (**Fig. 4j**). To test whether this activity-dependent omission reflected a specific CaImAn parameter choice, we performed a CaImAn parameter titration using the 3D nuclear masks as ground truth. At a fixed gSig = 8, CaImAn titration across selected min_corr and min_pnr combinations showed that relaxing min_pnr increased identity-level and active-neuron recall over an intermediate range, whereas overly permissive settings increased the false positive rate without further improving recall. Here, false positive rate was defined as the fraction of CaImAn components that failed to match a CLG-defined nucleus under the same 30-pixel cutoff. These results indicate that the omission was not explained by a single overly stringent parameter setting, but reflected a broader trade-off between neuronal recovery and unmatched components (**Supplementary Fig. 20**). Together, these findings show that the value of structural calibration in mouse cortex lies not only in consolidating duplicated detections, but also in reducing activity-dependent omission (**Supplementary Videos 14 and 15**).

Finally, we examined how these differences affect downstream network representations in mouse cortex. Using the same network analysis pipeline as for zebrafish, we reconstructed functional connectivity networks for both the full CLG-calibrated labeled population and the CaImAn-identified subset. Connection thresholds were normalized to the 90th percentile of pairwise correlations to account for size differences. In the full population, pre-and post-calibration networks differed significantly in degree and eigenvector centrality distributions (**Fig. 4h,i**; **K–S test, *P* < 0.0001** for both metrics), and calibration enhanced long-range activity correlations (**Fig. 4k**). Network properties from the full labeled population diverged markedly from those derived using CaImAn (**Fig. 4l,m**). Applying the adapted GDM dismantling algorithm ^40^, we found that node importance profiles and fragmentation dynamics varied significantly between the two network types (**Fig. 4n**). Notably, the most influential nodes identified in the CaImAn subset showed poor correspondence with those in the full network (**Fig. 4o**). Together, these results indicate that, in mouse cortex as in zebrafish, inferred network structure is distorted by both cross-plane duplication and activity-dependent omission. Structural calibration reduces these biases by anchoring functional signals to stable three-dimensional cellular identities.

## Discussion

This study defines single-neuron identity as a calibration problem in volumetric calcium imaging. In high-throughput recordings, one physical neuron can be represented in multiple axial planes and then converted by plane-wise segmentation into multiple functional nodes. This is not random noise. It is a structured identity error. In larval zebrafish, rapid functional imaging produced ∼80,000 plane-wise detections, whereas CLG calibration resolved these signals into ∼50,000 nucleus-defined single-neuron identities. In mouse visual cortex, the same failure mode persisted despite larger somata and lower cell density, with ∼40% of putative detections consolidated after calibration. These results show that the basic unit of analysis in volumetric calcium imaging—the node—cannot always be assumed to be a neuron.

The principle of CLG is simple: calcium activity reports function, but nuclear structure defines identity. By using nuclear labels as stable three-dimensional identity anchors, CLG assigns calcium signals to physical cells rather than to plane-wise fluorescent regions. This distinction matters most when axial sampling becomes coarse relative to the optical and anatomical extent of a soma. This situation is not limited to a single microscope design. It can occur in systems that intentionally elongate the effective axial sampling volume, such as underfilled-NA ^14,43^ or Bessel-beam ^44^ approaches, and also in high-throughput volumetric platforms that aim to capture more neurons, larger volumes or more axial planes within a short time window^45,46^. In such regimes, the push toward larger neuronal coverage can increase the risk of sampling the same cell multiple times and treating it as multiple functional nodes. Thus, the practical trade-off is not simply speed versus resolution, but throughput versus identity fidelity.

This distinction sets a clear limit on heuristic deduplication. Spatial-overlap, proximity, and correlation rules can remove some obvious cross-plane duplicates, but they infer identity from geometry and functional similarity rather than from physical structure ^22^. Our FACED-style benchmark shows the practical consequence. At the published threshold, the algorithm produced a –19.2% count error, split 50.2% of multi-plane nuclei across multiple clusters, and merged ROIs from different nuclei in 37.6% of clusters. Adjusting thresholds revealed a trade-off between false splits and false merges. A corrected cell count does not guarantee correct single-neuron identity.

CLG should therefore be viewed as an identity-calibration layer, not as a replacement for existing calcium-imaging pipelines. Calcium-only methods such as CaImAn infer neurons from activity patterns and are biased toward signals that are strong, isolated or temporally sparse^20^. In mouse visual cortex, CLG recovered all CaImAn-detected active neurons and additionally identified 2,032 active, nucleus-anchored neurons missed by CaImAn. Parameter titration showed that this was not explained by one overly stringent CaImAn setting; improving recall introduced a false-positive burden. Thus, structural calibration and calcium-signal extraction solve different problems. Existing denoising, demixing and spike-inference tools estimate activity. CLG defines which physical neuron that activity belongs to.

The same logic applies to network analysis. Graph-derived phenotypes amplify identity errors because the node definition is itself part of the measurement. When one neuron is represented as several ROIs, the graph is built from the wrong units. Calibration changed hub assignment^39^, communicability and robustness estimates^47,48^, showing that duplicated ROIs do not merely add noise; they can change the inferred circuit architecture.

The PTZ experiments demonstrate the practical consequence. After calibration, spontaneous and PTZ-treated networks showed consistent differences in long-range correlations, hub organization, and dismantling profiles. Without calibration, the same analysis produced unstable or contradictory results across animals. Thus, biological conclusions about hub reorganization and network resilience depended on whether functional nodes corresponded to true single-neuron identities.

CLG has limitations. It requires a structural reference channel, which adds genetic and optical complexity. Its performance depends on nuclear segmentation quality, structural–functional registration and residual contamination control during trace reassignment. The current implementation is best suited to soma-centered volumetric imaging, where nuclei provide reliable cell identity anchors. Sparse labeling, severe motion, chronic tissue deformation, low-SNR structural channels or analyses focused on dendrites and axons will require additional development. For datasets lacking nuclear labeling, optical parameters such as Z-step size, soma diameter and axial PSF width may provide a first-order estimate of duplication risk, but they cannot replace cell-level identity correction. Public large-scale calcium-imaging resources with cell-level segmentations or standardized ROI outputs may provide useful external test beds for future studies of identity errors across species, imaging modalities and analysis pipelines^49^.

Despite these limits, CLG provides a practical route to identity-consistent volumetric calcium imaging. It makes a hidden measurement variable explicit: before neural activity can be interpreted, functional signals must be assigned to stable physical cell identities. This requirement will become more important as volumetric imaging platforms increase speed, field of view and axial coverage. By anchoring functional signals to stable physical cell identities, CLG turns single-neuron identity from an unexamined assumption into a calibrated measurement.

## Methods

### Zebrafish husbandry and preparation

Wild-type AB and transgenic zebrafish (Danio rerio) lines were maintained under standard laboratory conditions at 28.5 °C on a 14 h light/10 h dark cycle in a recirculating aquatic system. All experimental procedures were performed in accordance with protocol IMM-XiongJW-3, approved by the Institutional Animal Care and Use Committee of Peking University and accredited by the Association for Assessment and Accreditation of Laboratory Animal Care International (AAALAC). Embryos and larvae were staged according to hours or days post-fertilization (hpf or dpf) and reared in E3 medium. To inhibit pigmentation and maintain optical transparency for imaging, 0.002% phenylthiourea (PTU; Sigma, St. Louis, MO) was added at 10 hpf. Only healthy, normally developed larvae were used for imaging experiments.

### Transgenic Zebrafish Generation

The Tg(*elavl3*:GCaMP6s) zebrafish line was obtained from Dr. Florian Engert at Harvard University. The Tg(*elavl3*:H2B-mRuby3) reporter zebrafish line was generated using Tol2 transposase RNA-mediated transgenesis, as previously described ^50^. The plasmid *elavl3*:H2B-mRuby3 construct was modified using the plasmid *elavl3:H2B-GCaMP6s* (available on Addgene) as a template. The *H2B-GCaMP6s* fragment was excised using the *Age*I-HF enzyme, and the backbone, H2B, and mRuby3 fragments were reconstituted using the ClonExpress MultiS One Step Cloning Kit (Vazyme). The final construct was co-injected with Tol2 transposase RNA into the eggs of wild-type AB strains to generate Tg(*elavl3*:H2B-mRuby3) F0 fish.

Primers used for plasmid construction:

- *h2b*-f: 5’ CCCAACCTGTTATATTTTCCACCTGC

- *h2b*-r: 5’ GAGATCCTTATCGTCATCGTC

-mRuby3-f: 5’ GACGATGACGATAAGGATCTCGCCACCATGGTGTCTAAGGG

- mRuby3-r: 5’ GTGGTTTGTCCAAACTCATC

### Surgical Procedures for Cranial Window Implantation and Viral Vector Delivery in Mice

All surgical and experimental procedures were approved by the Institutional Animal Care and Use Committee of Peking University. Male C57BL/6J mice (28–50 days old at surgery; 49–70 days old at imaging) were housed under standard conditions (12 h light/12 h dark cycle) with ad libitum access to food and water. To express fluorescent indicators, mice were injected with AAV-*hSyn*-GCaMP6s-P2A-H2B-mRuby3-*WPRE*-*hGH* polyA (5 × 10¹² vg/mL; injection rate 10 nL/min; volume 200 nL) 1–2 weeks before cranial window implantation. Bilateral injections targeted the visual cortex at AP: –1.58 mm, ML: ±0.75 mm, DV: –0.3 mm and AP: –2.46 mm, ML: ±0.75 mm, DV: –0.3 mm. During surgery, mice were anesthetized with isoflurane (1.0% maintenance, 0.5 L/min) and secured in a stereotaxic frame. After removing the scalp and cleaning the skull, a 4 mm diameter circular craniotomy was made over the target region, preserving the posterior 1–1.5 mm of bone to avoid major sinuses. A 4 mm glass coverslip was placed over the craniotomy and sealed with tissue adhesive, followed by 3M tissue glue and dental cement to secure the window. A titanium head plate was affixed to the skull, centered on the cranial window. Postoperative care included intraperitoneal injection of 1% ceftriaxone sodium (0.1 mg/10 g body weight) for three days. Mice were allowed at least one week of recovery before imaging. Animals with dural damage or opaque windows were excluded from experiments.

### Two-Photon Imaging of Zebrafish

For in vivo imaging, zebrafish larvae were embedded in low-melting-point agarose to mechanically stabilize the preparation during volumetric acquisition. This embedding procedure minimized large sample displacement and enabled reproducible alignment between functional volumes and the structural reference stack.

Imaging experiments were performed using an Olympus FVMPE-RS microscope equipped with a Spectra-Physics InSight X3 dual-output laser and a 25×/1.05 NA water-immersion objective. GCaMP6s and mRuby3 signals were simultaneously excited at 920 nm. GCaMP6s fluorescence was collected using a 495–540 nm emission filter, and mRuby3 fluorescence was collected using a 575– 630 nm emission filter.

Structural imaging was performed using the galvanometer scanning module at 920 nm excitation. Single-plane images were acquired at 1024 × 1024 pixel resolution (FOV: 510 × 510 µm, pixel size: 0.5 µm) with a 3 s acquisition time per plane. Z-axis scanning was performed at 1 µm intervals across a depth range of 200–300 µm.

Functional imaging was performed using the resonant scanning module at 920 nm to capture rapid volumetric dynamics. Volumes comprising 27–30 slices were acquired at a volumetric rate of 1 Hz, with 5–7 µm slice intervals, covering a depth range of ∼200 µm. For pharmacological interventions, seizure-like symptoms were induced by treating zebrafish with 10 mM pentylenetetrazole (PTZ) for 30 min, and brain activity was recorded post-treatment.

### Two-Photon Imaging of Mice

Awake mice were head-fixed in an imaging holder using screws and secured on the imaging platform. Imaging was performed using the same Olympus FVMPE-RS microscope and 25×/1.05 NA water-immersion objective described above. Unless otherwise stated, optical settings (including excitation wavelength and emission filters) were consistent with the zebrafish experiments.

Functional imaging of GCaMP6s signals was performed using the resonant scanning module at 920 nm. Volumetric time-series were acquired at a scanning speed of 2.36 Hz with 20 µm axial intervals across a depth range of 0–260 µm. Corresponding high-resolution structural images were acquired at 2 µm Z-steps to provide the 3D nuclear masks for segmentation.

### Image Preprocessing

Image preprocessing encompassed both structural and functional datasets. For mRuby3 structural images, preprocessing included denoising and local normalization to enhance single-nucleus identification accuracy. We used the sparse deconvolution algorithm (publicly available MATLAB package)^24^ in this paper: Parameters included 120 sparse iterations, Z-axis continuity of 1, image fidelity of 150, sparsity of 6, and 8 deconvolution iterations.

Local normalization was applied to address uneven illumination and enhance foreground-background contrast. The sliding window local normalization formula is as follows:

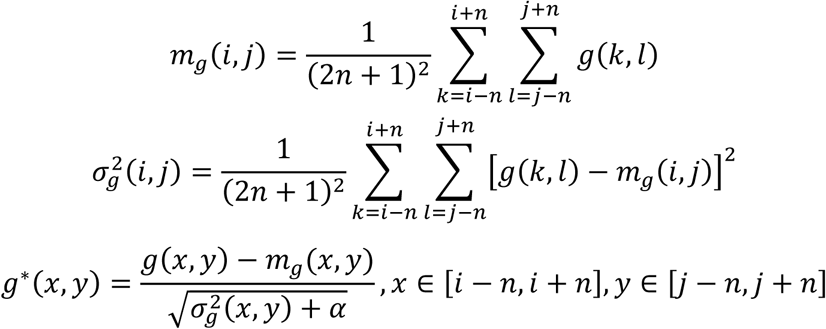

Here, *m*_g_(*i*, *j*) and σ^2^_*g*_(*i, j*) represent the mean and variance of pixels within a radius *n* centered at (*i*, *j*), respectively. *g*^∗^(*x*, *y*) is the normalized pixel value at (*x*, *y*), and α is a regularization parameter (set to 1 × 10^−3^). The radius n was set to be larger than the diameter of the nucleus in pixels. In this study, n was chosen to range between 10–30 pixels, ensuring that the sliding window adequately covered the nucleus size for accurate local normalization.

For functional images, GCaMP6s signals were denoised using the SUPPORT self-supervised DL algorithm ^51^. A blind-spot network was trained for 20 epochs on a desktop equipped with an NVIDIA 3080 GPU. The model was trained on a single time-series functional imaging result and could be applied to denoise other datasets of the same type, ensuring consistent noise reduction across experiments.

### Functional Image Registration

To align the rapid volumetric functional data (512 × 512 pixels) with the high-resolution structural reference (1024 × 1024 pixels), functional images were first spatially upsampled to match the structural resolution (1024 × 1024 pixels). Motion correction and non-rigid registration were subsequently performed using the NoRMCorre algorithm ^52^. Accordingly, the registration parameters were scaled to match the upsampled resolution, using a patch size of 256 pixels and an overlap of 64 pixels (equivalent to 128 and 32 pixels at the native 512 × 512 resolution, respectively) with 2 iterations.

Registration was driven by the GCaMP6s channel to ensure precise alignment between functional dynamics and the structural nuclear map. Specifically, the upsampled functional GCaMP6s time-series was registered to the high-resolution structural GCaMP6s image. Although the structural GCaMP6s image represents a slow-acquisition static snapshot, it preserved sufficient stable anatomical landmarks—including neuropil architecture, nerve fiber bundles, and high-contrast structural corners—to serve as a robust registration template. Because imaging was performed under mechanically stabilized preparations, including head-fixed mice and agarose-embedded zebrafish larvae, large axial displacement during acquisition was limited. Accordingly, the nominal Z-step recorded by the microscope provided the primary basis for assigning each functional plane to the structural volume. An optional manual inspection step was available to verify local plane-to-structure correspondence using anatomical landmarks and to make minor refinements when needed, but this was implemented as an auxiliary improvement rather than a required part of the alignment procedure. This strategy ensured that the motion-corrected functional data were spatially aligned with the 3D nuclear coordinates defined by the structural mRuby3 channel.

### Nucleus Segmentation

A training dataset of over 10,000 zebrafish larval nuclei was manually annotated in QuPath, along with a smaller test dataset for algorithm evaluation. These datasets were used to train and benchmark Cellpose 2.0, StarDist, and Mask R-CNN. Cellpose 2.0 and StarDist performed best, and Cellpose 2.0 was selected for final 3D nucleus segmentation in mRuby3 structural images.

For mouse brain regions, where nuclei are more sparsely distributed, we utilized the default pre-trained model bundled with the StarDist ImageJ plugin for initial nucleus detection. Remaining annotation errors were manually corrected in the Cellpose 2.0 GUI, where StarDist predictions were refined. The corrected dataset was then used to train a customized Cellpose 2.0 model, which was subsequently applied to other mouse brain region images.

### Quantification of Nuclear Size

Morphological properties of segmented nuclei were analyzed using the regionprops3 algorithm (MATLAB). To account for the irregular shapes of neuronal nuclei in 3D space, nuclear size was quantified as the equivalent spherical diameter, defined as the diameter of a sphere with the same volume as the segmented object. This metric is calculated as:

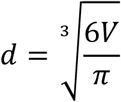

where *V* represents the physical volume of the nuclear mask, derived by multiplying the total number of voxels within the segmented region by the voxel dimensions. This approach provides a standardized geometric measure for comparing neuronal sizes across different brain regions and developmental stages.

### Single-Neuron Signal Extraction

The mRuby3 structural images provided 3D nuclear masks for single-neuron assignment. The plane-to-structure correspondence established in the registration step was used to identify the 2D nuclear masks associated with each GCaMP6s functional plane. To assess the robustness of this assignment against residual axial uncertainty, we performed a sensitivity analysis: perturbing the Z-alignment of 10% of functional planes by 1–2 structural Z-planes (deviating from the optimal structural match) changed the total cell count by less than 0.2% (estimated across 100 randomized iterations), whereas perturbing 50% of planes changed it by only ∼1%. These results indicate that the final assignments were robust to minor alignment uncertainty. This procedure yielded colocalized 2D nuclear masks corresponding to the functional images. To enhance the SNR, ROIs with areas below four pixels were excluded. Time-series GCaMP6s signals were then extracted by averaging fluorescence intensity across pixels within each ROI, representing single-neuron activity at the imaged depth. Because each 3D nuclear mask carried a unique identifier, signals from different functional axial planes corresponding to the same neuron were merged by averaging their activity traces. Single-neuron ΔF/F signals were computed from GCaMP6s fluorescence using the allensdk.brain_observatory.dff module from the AllenSDK.

The signal extraction pipeline for mice was identical to that described for zebrafish. We processed the imaging results using our method and compared the extracted single-neuron signals with those obtained using CaImAn ^20^. To assess whether the activity-dependent omission observed in CaImAn could be mitigated by parameter adjustment, we additionally performed a CaImAn parameter titration on the mouse visual cortex dataset. CaImAn was rerun across 20 selected initialization-parameter combinations while keeping the remaining processing steps fixed. The spatial scale parameter was fixed at gSig = 8. The tested combinations were: min_corr = 0.60 with min_pnr = 3, 5, 7 and 10; min_corr = 0.70 with min_pnr = 3, 5, 7 and 10; min_corr = 0.80 with min_pnr = 3, 5, 7, 10, 20 and 30; min_corr = 0.85 with min_pnr = 10, 20 and 30; and min_corr = 0.90 with min_pnr = 10, 20 and 30. For each run, CaImAn spatial components were matched to the nearest nucleus in the corresponding structural plane using the centroid of the spatial footprint, and detections within 30 pixels were counted as ground-truth matches. Identity recall was defined as the fraction of unique nucleus identities recovered after CLG calibration. Active and silent nuclei were classified using the same ΔF/F threshold of 0.13 as in the main analysis. The false positive rate was defined as the fraction of CaImAn components whose nearest nucleus exceeded the 30-pixel matching cutoff. This analysis enabled us to quantify the trade-off between neuronal recovery and false-positive burden across CaImAn parameter settings.

### FACED-style ROI-chain clustering benchmark

To evaluate whether proximity-and activity-based cross-plane ROI clustering could recover CLG-defined single-neuron identities, we implemented a FACED-style ROI-chain clustering benchmark based on the published FACED2 analysis description. Plane-wise ROI traces were taken from the pre-calibration ROI outputs. The original 3D CLG/Cellpose label mask and the functional-to-structural slice mapping were used to reconstruct the ROI footprint, centroid position and extraction order for each plane-wise ROI. The reconstructed ROI order was validated against exported nucleus identity and position arrays before benchmarking. CLG nucleus identities were withheld during clustering and used only for post hoc evaluation.

Adjacent-plane ROI chains were constructed using XY mask overlap between ROI footprints in neighboring functional planes. For each ROI, all non-zero overlapping labels in the adjacent plane were considered candidate links. When one ROI overlapped multiple candidates, the candidate with the highest Pearson correlation between calcium traces was retained, following the FACED-style ambiguity-resolution rule. Each connected component of these adjacent-plane overlap links defined a candidate ROI chain.

Within each ROI chain, ROIs were clustered using complete-linkage constraints. Each ROI was initialized as a singleton cluster. Candidate ROI pairs within a chain were ranked by trace Pearson correlation and considered for merging if their 3D centroid distance was below a distance threshold and their trace correlation exceeded a correlation threshold. Before accepting a merge, all ROI pairs in the proposed merged cluster were required to satisfy both criteria. Singleton ROIs not merged with any other ROI were retained as predicted clusters.

We evaluated the published zebrafish FACED-style operating point, defined as 3D distance < 20 µm and Pearson correlation > 0.5, and performed a threshold sweep over distance cutoffs of 8, 10, 12, 15, 20, 25 and 30 µm and Pearson correlation cutoffs of 0.3, 0.4, 0.5, 0.6, 0.7, 0.8, 0.85, 0.9 and 0.95. Because the structural z-step was 1 µm and the functional-to-structural mapping table encoded the corrected functional depth, 3D distances were computed in micrometers using x–y pixel sizes of 0.5 µm and mapped structural z positions in 1-µm units.

Clustering performance was evaluated against withheld CLG nucleus identities. Count error was defined as (*N*_FACED_ — *N*_CLG_)/*N*_CLG_, where *N*_FACED_ is the number of predicted clusters including singleton ROIs and *N*_CLG_ is the number of unique CLG nucleus identities. False split rate was computed among CLG nuclei represented in multiple functional planes as the fraction whose ROIs were assigned to more than one predicted cluster. Cluster-level false merge rate was defined as the fraction of predicted clusters containing ROIs from more than one CLG nucleus identity. ROI-weighted false merge rate was defined as the fraction of all ROIs assigned to such mixed-identity predicted clusters.

For pair-level metrics, we constructed a local adjacent-plane pair universe using ROI pairs from neighboring functional planes within a 25-µm 3D radius. A pair was treated as ground-truth positive if the two ROIs shared the same CLG nucleus identity, and as predicted positive if the two ROIs were assigned to the same FACED-style predicted cluster. Pair precision, recall and F1 were computed from the resulting true-positive, false-positive and false-negative pair counts. Pair-distribution plots used all same-nucleus pairs and a downsampled set of different-nucleus pairs for display; the full local pair universe was used for metric calculation.

## Data Processing and Analysis

### Network analysis of zebrafish whole-brain neuronal activity

Single-cell calcium activity traces were further processed to reduce noise and improve signal fidelity prior to network construction. Raw fluorescence images were first denoised using the SUPPORT self-supervised deep learning algorithm (see *Image Preprocessing*). Following denoising and computation of Δ*F*/*F* signals (see *Single-Neuron Signal Extraction*), the resulting single-cell activity matrix (N×T, where N is the number of neurons and T is the number of frames) was subjected to principal component analysis (PCA) to suppress observational noise. Principal components collectively explaining 95% of the total variance were retained, and neuronal activity traces were reconstructed from these components to enhance the SNR.

Functional connectivity networks were then inferred by computing pairwise Pearson correlation coefficients between reconstructed neuronal activity traces. Edges were defined for absolute correlation values exceeding 0.95 to sparsify the network and obtain a conservative estimate of functional coupling, thereby minimizing spurious correlations in high-density neuronal datasets. Given the large scale of the reconstructed whole-brain networks (N > 50,000 nodes), computing advanced topological metrics using standard libraries (e.g., NetworkX) proved computationally prohibitive. Therefore, we implemented accelerated algorithms for key metrics:

1. Eigenvector Centrality: We computed the leading eigenvector of the adjacency matrix using the Lanczos algorithm (scipy.sparse.linalg.eigsh), which efficiently retrieves the principal eigenvalue and eigenvector without full matrix decomposition. To enable fair comparison between pre-and post-calibration networks with differing node counts (*N*_pre_ vs. *N*_post_), centrality scores were normalized by the *L*_1_ norm and scaled by a factor of *N*_pre_/*N*_post_ to account for the reduction in network size due to CLG calibration.
2. Communicability: Instead of full matrix exponentiation, we adopted the definition and efficient computation strategy described by Munn et al. (2024) ^39^, which quantifies the weighted sum of walks between nodes, capturing multiscale communication efficiency across the network.

Other basic metrics (degree, clustering coefficient) were computed using NetworkX v3.2.1(Python 3.11).

### Statistical Analysis of Zebrafish Whole-Brain Neuronal Activity

The correlation function *g*(*r*) was calculated as follows:

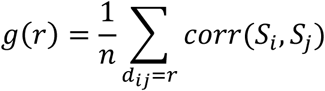

Here, *d*_ij_ is the Euclidean distance between neurons *i* and *j*, *n* is the number of neuron pairs with *d*_ij_ = *r*, and *corr*(*S*_i_, *S*_j_) is the Pearson correlation coefficient between their activity signals *S*_i_ and *S*_j_. In this study, we computed *g*(*r*) for all neuron pairs across the zebrafish brain and generated a histogram by binning the results based on Euclidean distance r.

### Network dismantling analysis

To assess network robustness and identify critical nodes, we performed a network dismantling analysis. Given the computational intractability of applying dismantling algorithms directly to full single-cell networks (*N* > 50,000), we first applied a hierarchical coarse-graining procedure following the geometric renormalization framework described by Munn et al. ^39^. Briefly, neurons were iteratively paired and merged based on the maximal pairwise Pearson correlation of their temporal fluorescence traces. In each step, highly correlated pairs were fused into a single ’super-node,’ and their activity signals were averaged to represent the new unit. This recursive 2-to-1 mapping generated a multiscale representation of the network. For subsequent analyses, we selected a coarse-graining level where each super-node represents the aggregate activity of 256 original neurons, providing an optimal balance between topological complexity and computational efficiency.

Network dismantling was then performed on the coarse-grained networks using two complementary methods. Our primary method was a modified version of the Graph Dismantling with Machine Learning (GDM) algorithm developed by Grassia et al. ^40^. To make the training examples more representative of neural-network topology, we supplemented the original training set with synthetic Watts–Strogatz and modular graphs containing 25 nodes each, capturing high clustering and modular organization characteristic of zebrafish brain networks. To validate the performance of our modified algorithm, we also applied the CoreHD dismantling method ^41^ for comparison. The adapted GDM algorithm not only achieved more efficient dismantling but also exhibited enhanced sensitivity in distinguishing topological differences between spontaneous and PTZ-induced epileptic states.

### Neuronal Avalanche Analysis

Spatiotemporal propagation of neuronal activity was assessed using neuronal avalanche analysis, adapting the framework described by Burrows et al. ^18^, with code obtained from their public repository (https://github.com/dmnburrows/criticality).

Signal Deconvolution and Binarization: To infer underlying spiking activity, raw fluorescence signals were processed using the CaImAn-MATLAB toolbox (https://github.com/flatironinstitute/CaImAn-MATLAB). We employed the thresholded OASIS algorithm with a first-order autoregressive model. To ensure solution sparsity and suppress noise, the regularization parameter was set to *λ* = 10, combined with a noise-dependent constraint (*s*_min_>3*σ*). Subsequently, the deconvolved traces were binarized using a fixed amplitude threshold to generate a discrete event matrix representing the timing of neuronal activations.

Avalanche Definition and Parameter Estimation: The temporal resolution was determined by the image acquisition rate, resulting in a time bin width of Δ*t* = 1 s. A neuronal avalanche was defined as a contiguous sequence of time bins containing at least one active neuron, bounded by inactive bins. For each avalanche, we quantified its size (*S*) (total number of binary events) and duration (*D*) (number of consecutive active bins). The power-law exponents for avalanche size (*τ*) and duration (*α*) distributions were estimated using the computational tools provided by Burrows et al.

## Code Availability

The custom software suite for Comprehensive Label-Guided (CLG) framework, including modules for image preprocessing, 3D structural segmentation, calibration, and network analysis, is available on GitHub at https://github.com/PKUCHENLAB/CLG-Volumetric-Imaging-Analysis-Framework/. A version of the code used to generate the results in this manuscript is also provided as Supplementary Software. The repository also includes the scripts for reproducing the network dismantling. We utilized the following open-source algorithms: Sparse Deconvolution (https://github.com/WeisongZhao/Sparse-deconvolution) for structural image resolution enhancement, Cellpose 2.0 (https://github.com/MouseLand/cellpose) for segmentation, NoRMCorre (https://github.com/flatironinstitute/NoRMCorre) for motion correction, and the SUPPORT algorithm for denoising. The modified GDM network dismantling algorithm is included in our GitHub repository. A persistent DOI for the code version used in this publication will be made available upon acceptance.

## Data Availability

The DL model weights trained on zebrafish and mouse datasets, along with the manually annotated dataset of >10,000 nuclei used for training, will be available at Zenodo under accession code 10.5281/zenodo.17775377. Due to the large size of the raw volumetric functional imaging datasets (which exceeds standard repository limits), the full raw data are available from the corresponding authors upon reasonable request. To facilitate reproducibility, we have provided a representative sample dataset (including both structural and functional channels) at Zenodo. Extracted single-neuron activity traces and 3D spatial coordinates underlying the analyses in Figures 1–4 and Supplementary

Figures are provided as Source Data.

## Author Contributions

L.C., C.G. and H.L. conceived the project. L.C., C.G., J.W. and L.M. supervised the study. X.L. and D.G. designed the research, developed the CLG analysis framework, and performed data analysis under the supervision of L.C. and C.G. D.G. prepared samples, performed two-photon imaging experiments on zebrafish and mice, and assisted with data processing. D.G. and C.S. developed image analysis methods, performed cell segmentation, manually annotated ground truth datasets, and optimized the segmentation pipeline. X.L. and J.Z. constructed and analyzed functional networks and performed graph dismantling analysis. X.L. composed the figures and videos under the supervision of L.C. S.R., L.X. and Y.L. (Yanmei Liu) assisted with sample preparation. Y.L. (Ye Liang) and H.M. assisted with optical experiments and equipment debugging. M.L. and X.L. performed 3D data visualization under the supervision of L.M. L.C. and X.L. wrote the manuscript with input from all authors. All authors discussed the results and commented on the manuscript.

## Acknowledgements

We thank Dr. Florian Engert for providing the transgenic zebrafish lines. We acknowledge Shaoling Qi (Olympus/Evident China Life Science) for technical support with the microscopy system. We are grateful to Jingzhong Lu for valuable advice on image processing, and to Wenzheng Wei (Hangzhou Chizha Technology Co., Ltd.) and Haocheng Long for software technical support. This work was supported by the National Natural Science Foundation of China (grants T2288102, 32227802 and 82525105 to L.C.), the Scientific Research Innovation Capability Support Project for Young Faculty (ZYGXQNJSKYCXNLZCXM-H7 to H. L.), the National Key Research and Development Program of China (grant 2022YFC3400600 to L.C.), the Collaborative Research Fund of the Hong Kong Research Grants Council (grant C7015-23G to J.W.), and the State Key Laboratory of Advanced Manufacturing for Optical Systems (grant KLMSKF202402 to C.G.). This work is supported by Biomedical Computing Platform of National Biomedical Imaging Center, Peking University.

