## Supplementary Information for "Structural–functional calibration corrects single-neuron identity errors in volumetric calcium imaging"

### Table of Contents

1. **Supplementary Figures and Legends (S1 – S20)**
2. **Supplementary Video Legends**

### 1. Supplementary Figures and Legends

CaImAn

Cite-On

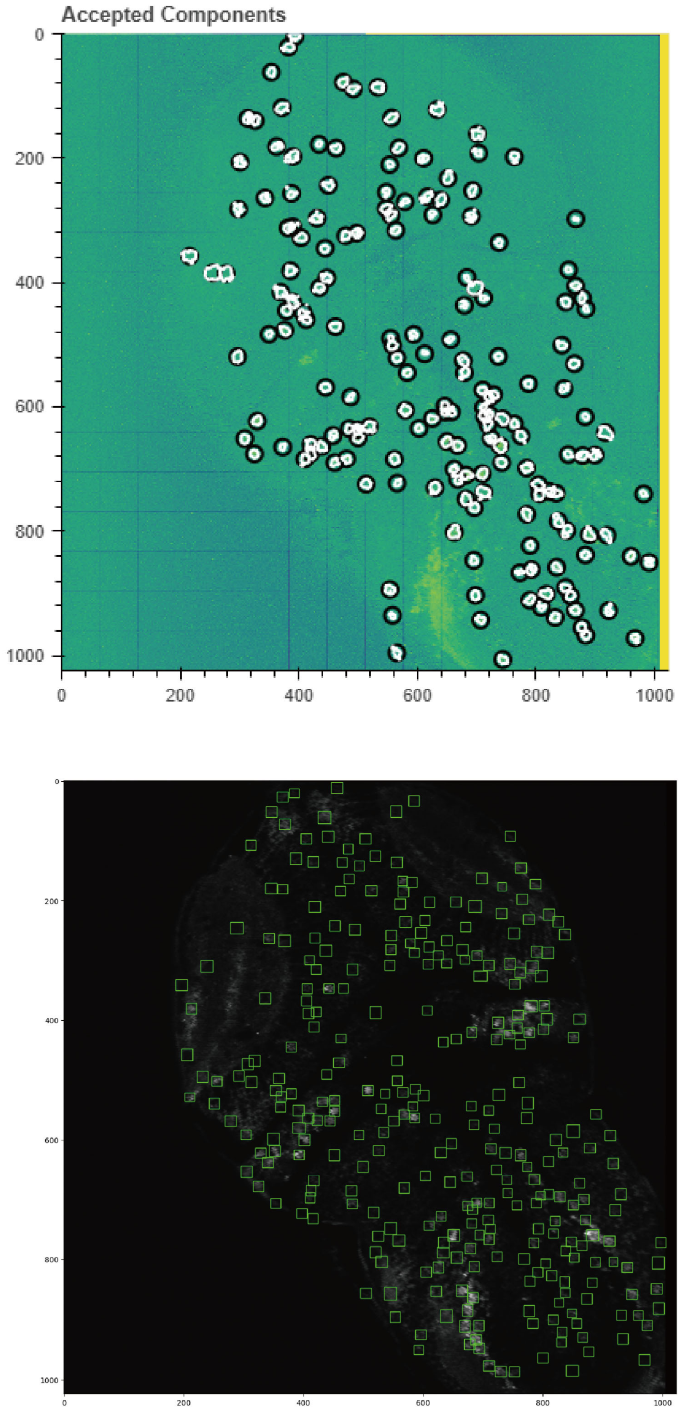


**Supplementary Figure 1. Segmentation results using calcium signal–based neuron extraction methods.**

Comparison of conventional approaches (e.g., non-negative matrix factorization in CaImAn) and deep learning–based algorithms (CITE-on) for neuron extraction in zebrafish.

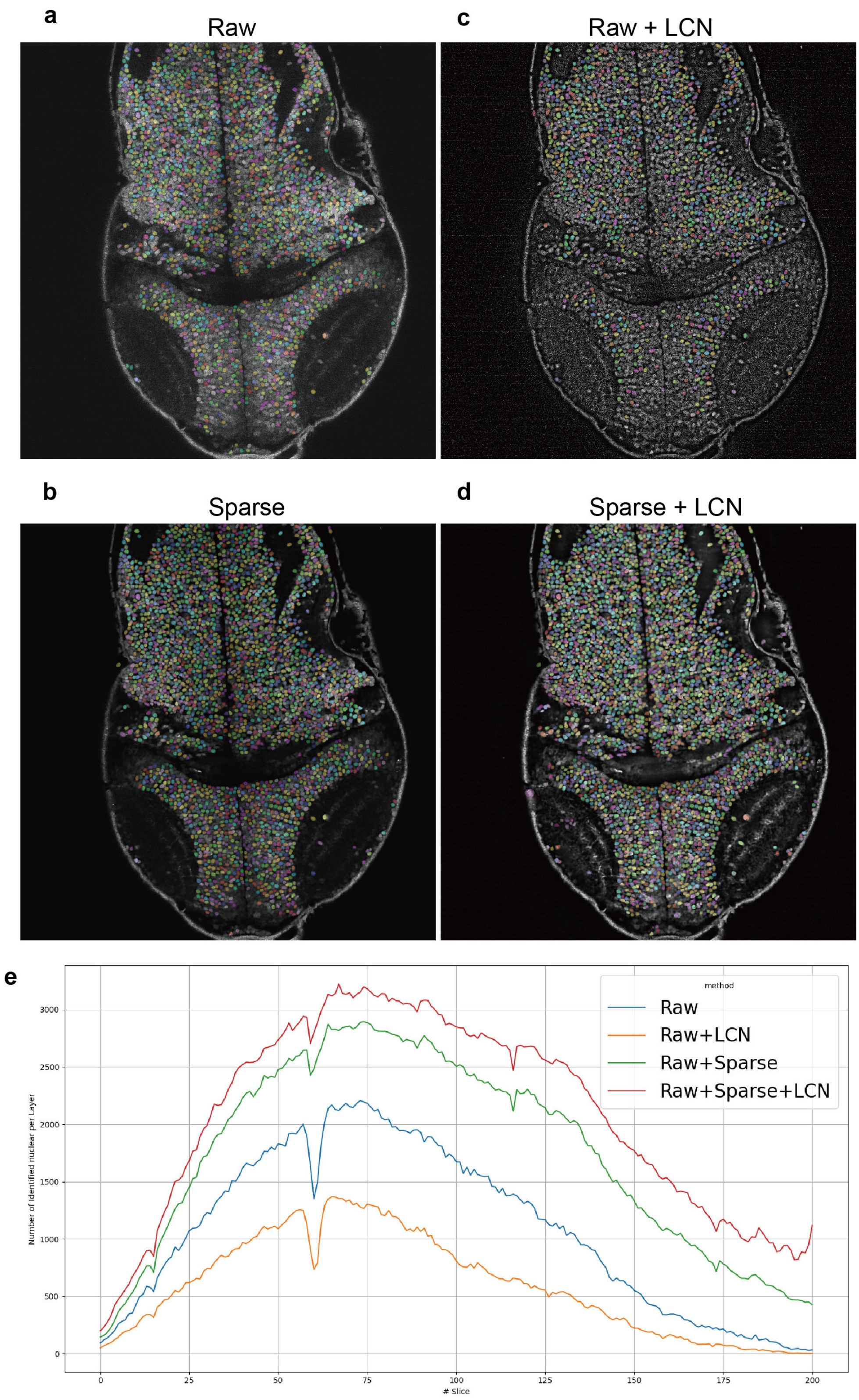


**Supplementary Figure 2. Image quality improvement by sparse deconvolution and local normalization.**

**a–d,** Segmentation results under four processing conditions. **e**, Cell counts per imaging layer across brain depths.

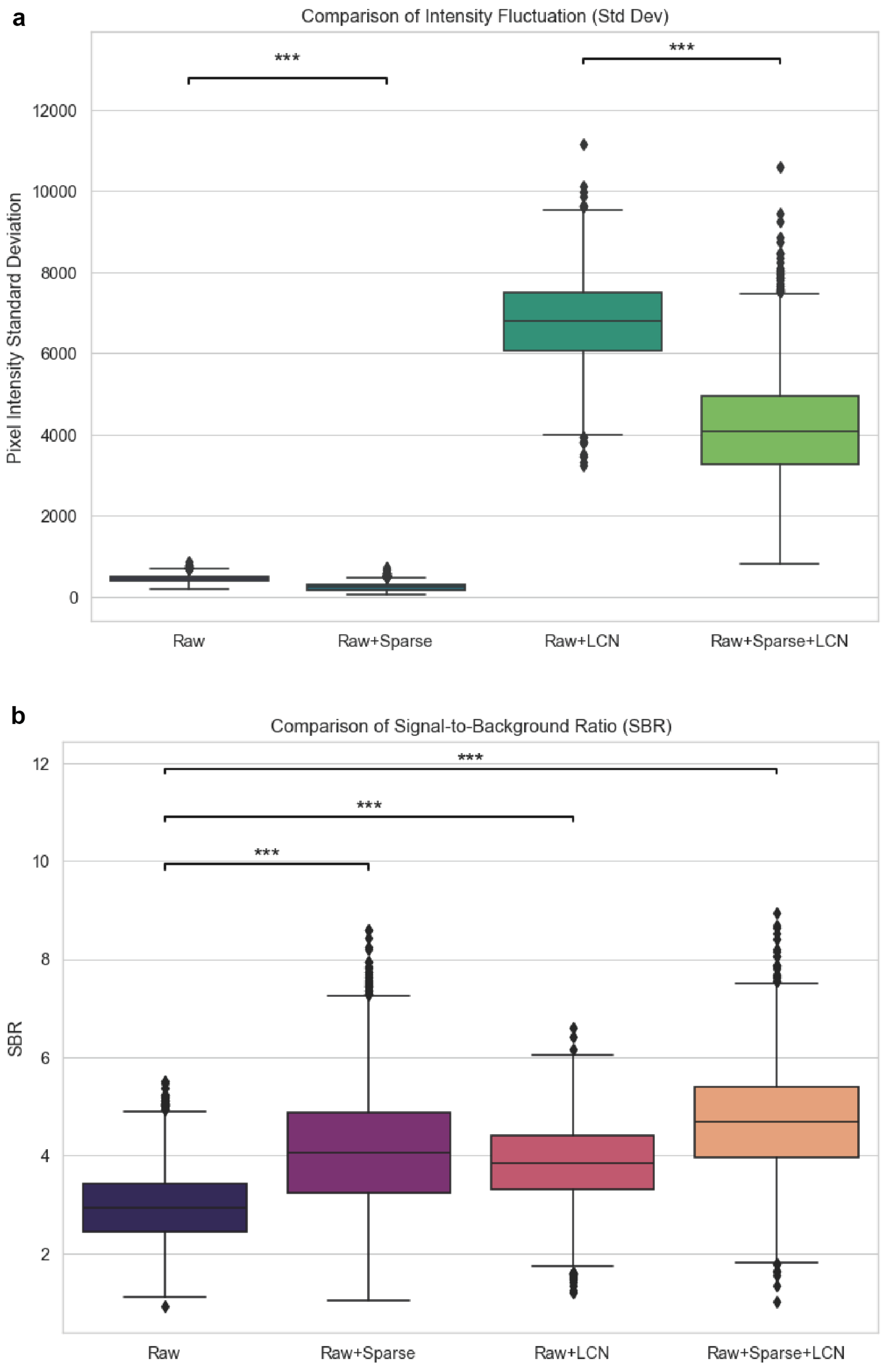


**Supplementary Figure 3. Quantitative evaluation of preprocessing effects.**

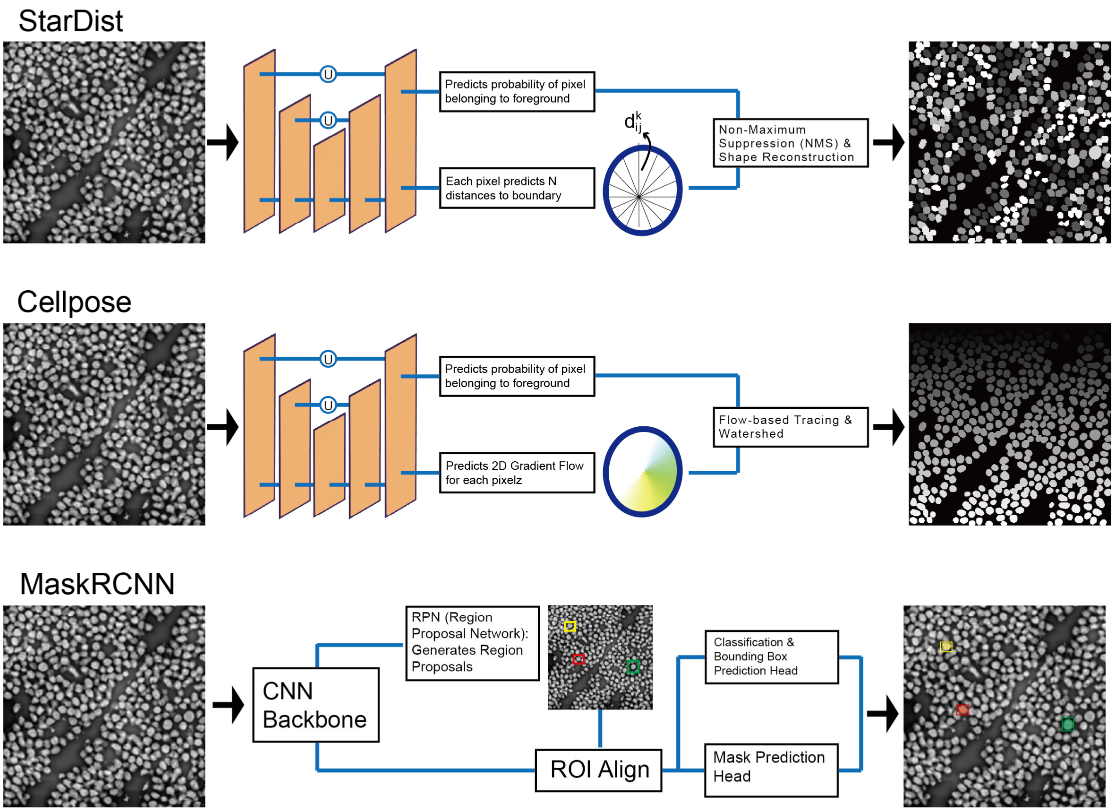


**Supplementary Figure 4. Deep learning–based segmentation algorithms.**

Schematic comparison of StarDist, Cellpose, and Mask R-CNN.

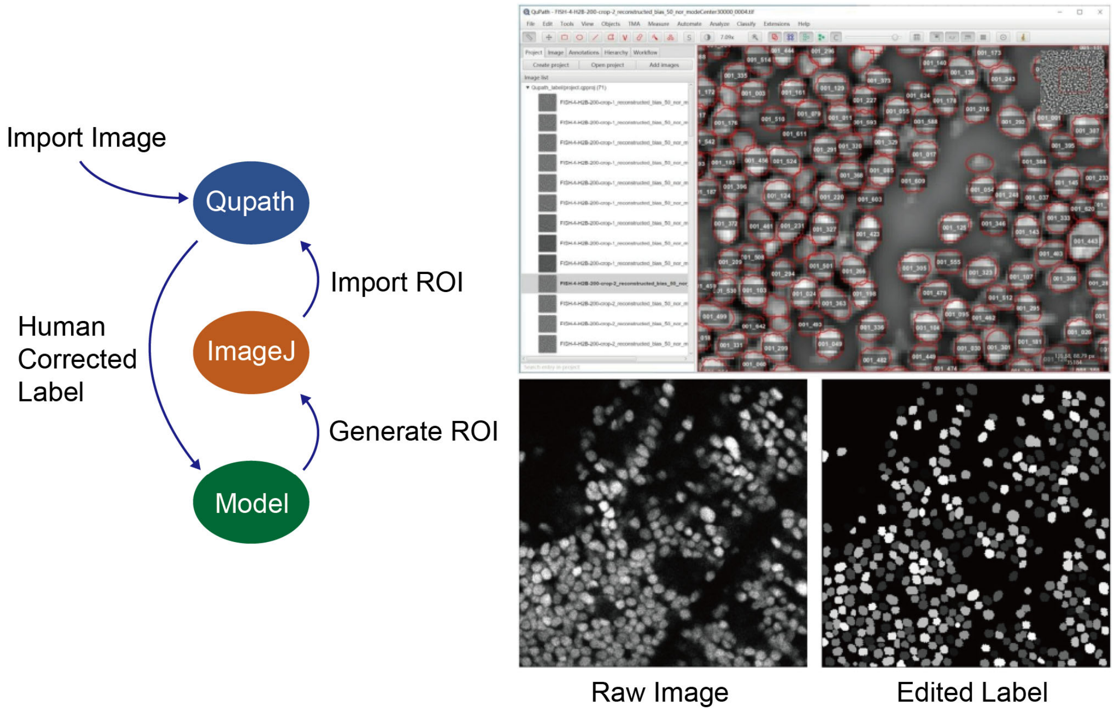


**Supplementary Figure 5. Ground-truth dataset generation.**

Workflow for manual annotation of >10,000 nuclei across diverse brain regions and imaging conditions.

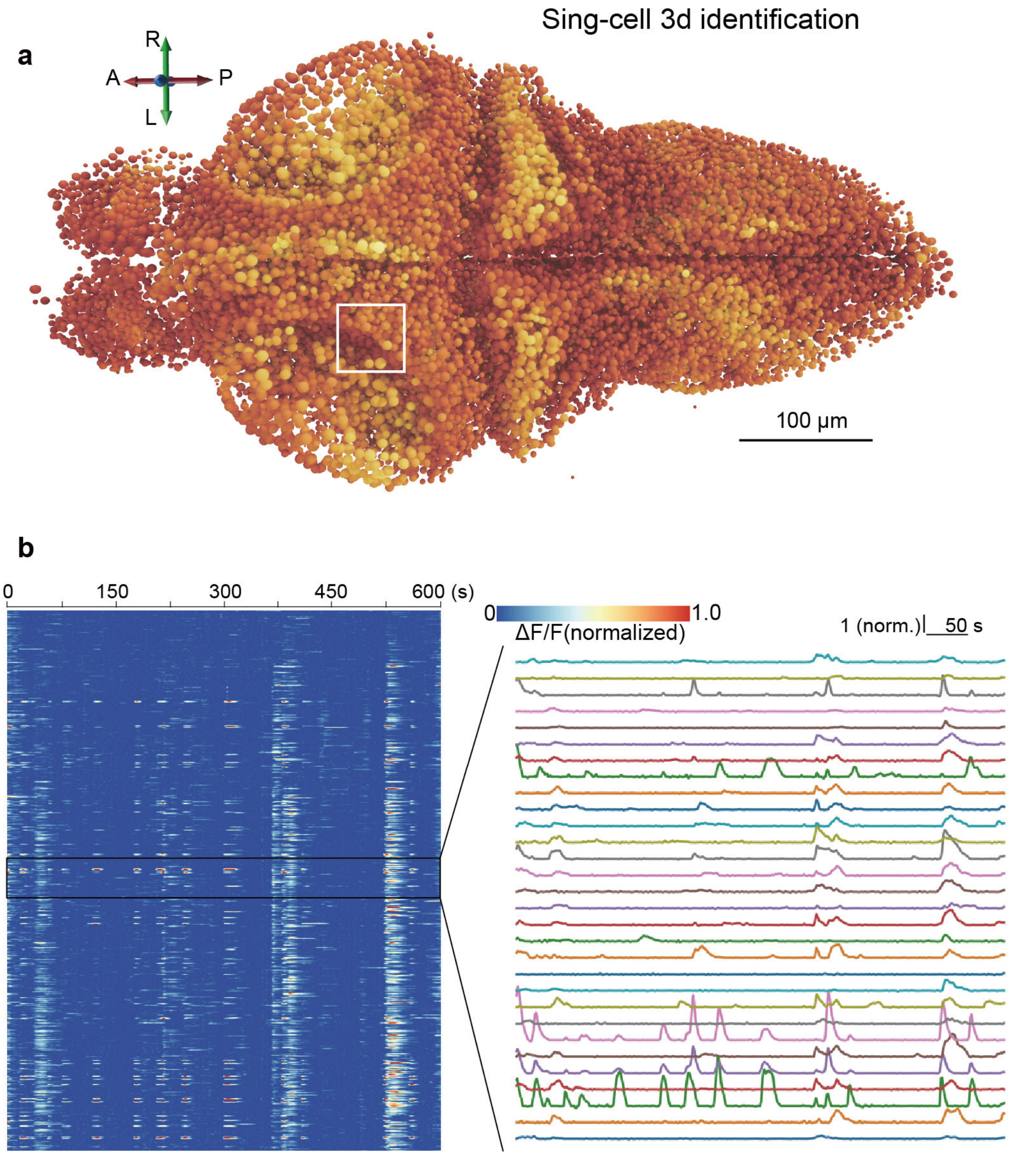


**Supplementary Figure 6. Accurate single-cell $Ca^{2+}$ trace extraction.**

**a,** 3D reconstruction of identified nuclei along the dorsal–ventral axis. **b,** Heatmaps of neuronal activity over 600 s, with magnified views highlighting single-cell dynamics.

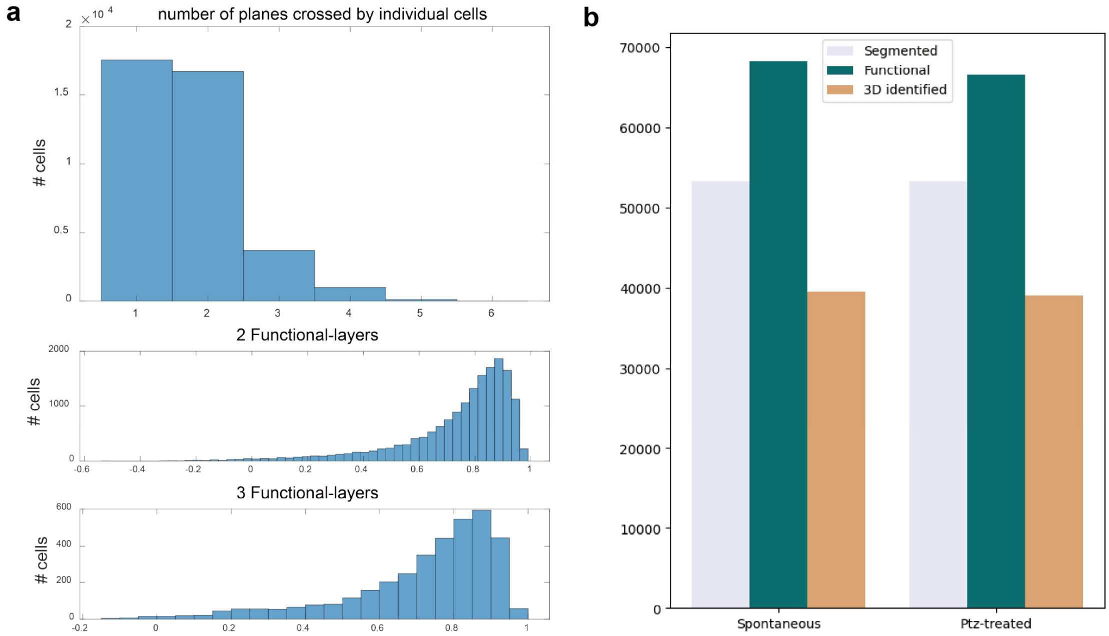


**Supplementary Figure 7. Neurons spanning multiple imaging planes.**

**a,** Distribution of neurons spanning multiple functional planes in PTZ experiments and pairwise correlations between signals from different planes. **b,** Comparison of cell counts from structural segmentation, functional imaging planes, and CLG-calibrated identification under PTZ-treated and spontaneous conditions.

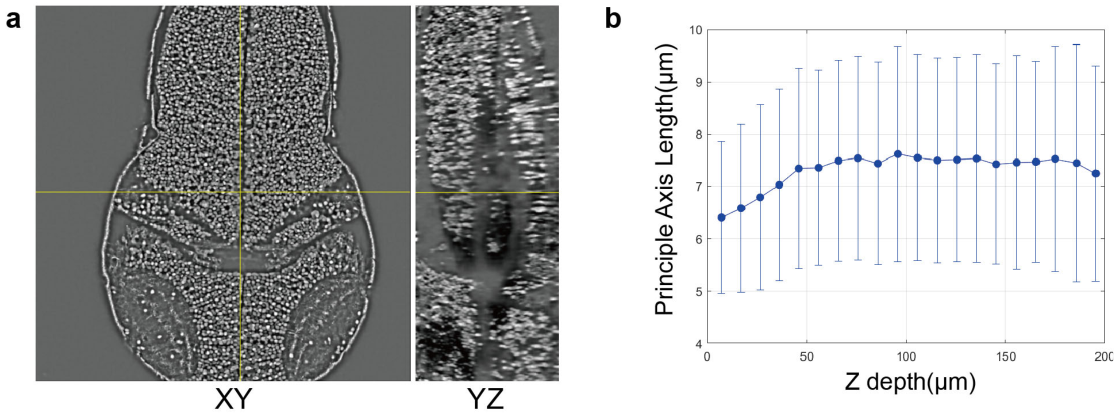


**Supplementary Figure 8. Depth-dependent degradation of axial resolution.**

**a,** XY and YZ cross-sections of structural images. **b,** Principal axis length of 3D-segmented nuclei as a function of z-depth.

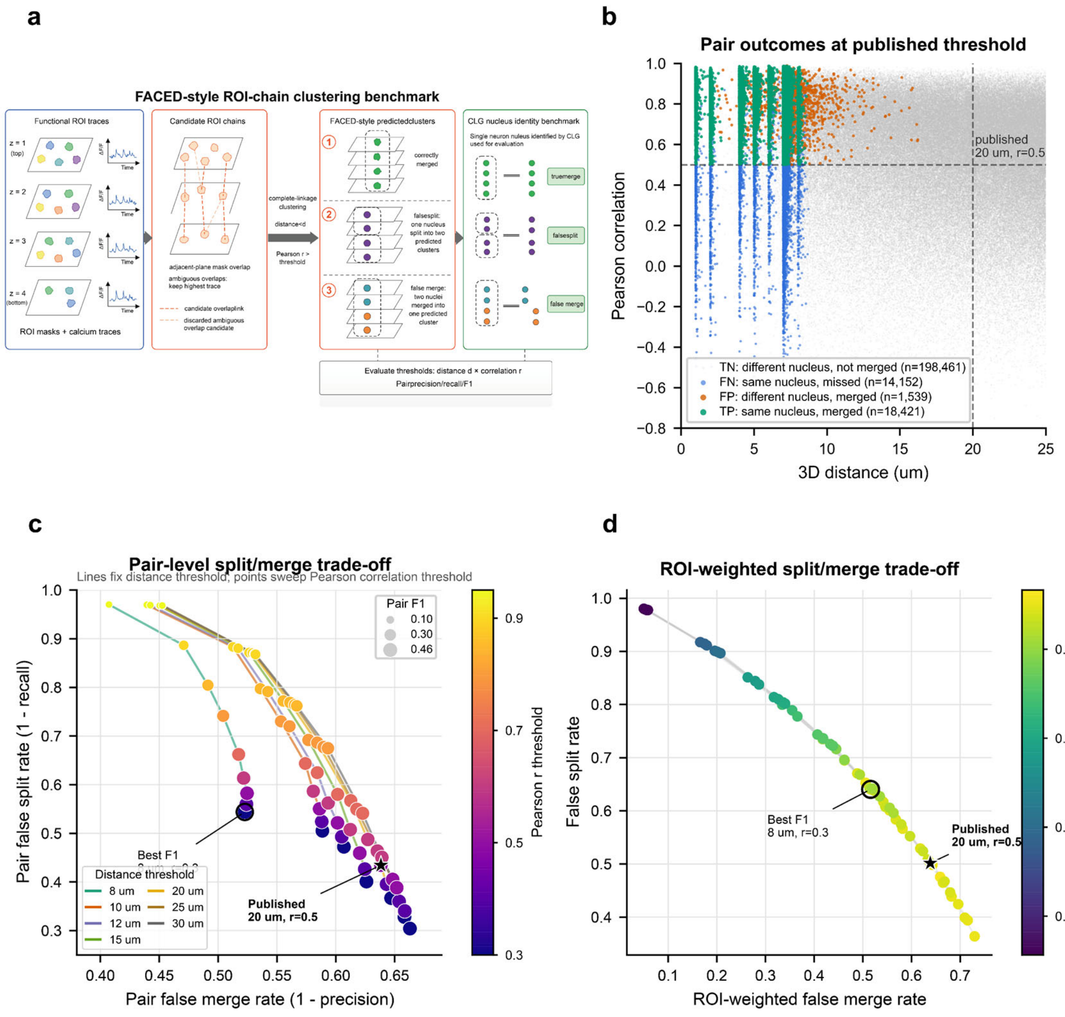

**Supplementary Figure 9. FACED-style ROI-chain clustering benchmark against CLG-defined nucleus identities.**

a, Workflow schematic. Plane-wise ROI masks and calcium traces were used as clustering inputs. Adjacent-plane candidate ROI chains were first constructed from XY mask overlap, with ambiguous overlap candidates resolved by retaining the candidate with the highest trace Pearson correlation. ROIs within each chain were then clustered using complete-linkage constraints requiring all ROI pairs within a predicted cluster to satisfy both a 3D-distance threshold and a trace-correlation threshold. CLG nuclear identity labels were withheld during clustering and used only for evaluation.

b, Local adjacent-plane ROI-pair outcomes at the published FACED-style zebrafish threshold. Displayed ROI pairs are plotted by 3D distance and calcium-trace Pearson correlation. Dashed lines mark the published operating point, 3D distance < 20 μm and Pearson r > 0.5. Colors indicate pair-level outcomes relative to CLG nucleus identity labels: true positives are same-nucleus pairs merged by FACED-style clustering; false negatives are same-nucleus pairs missed by clustering; false positives are different-nucleus pairs incorrectly merged; and true negatives are different-nucleus pairs not merged. Different-nucleus pairs were downsampled for visualization, whereas full local-pair counts were used for metric calculation.

c, Pair-level split/merge trade-off across threshold sweeps. The x axis shows pair false merge rate, defined as 1 − pair precision, and the y axis shows pair false split rate, defined as 1 − pair recall. Lines connect points with the same 3D-distance threshold while sweeping the Pearson correlation threshold. Point color encodes the correlation threshold and point size encodes pair-level F1. The star denotes the published FACED-style threshold and the open circle marks the best-F1 setting.

d, ROI-weighted and nucleus-level split/merge trade-off. The x axis shows the fraction of ROIs assigned to mixed-identity predicted clusters, and the y axis shows the fraction of multi-plane CLG nuclei split across multiple predicted clusters. Point color encodes pair-level F1. Across thresholds, relaxed clustering reduced false splits at the cost of increased false merges, indicating that proximity-correlation ROI-chain clustering removes some duplicate ROIs but does not fully recover CLG-defined single-neuron identities in this dataset. At the published threshold, this benchmark yielded a 19.2% undercount, a 50.2% false split rate among multi-plane nuclei, a 37.6% cluster-level false merge rate and a 63.9% ROI-weighted false merge rate.

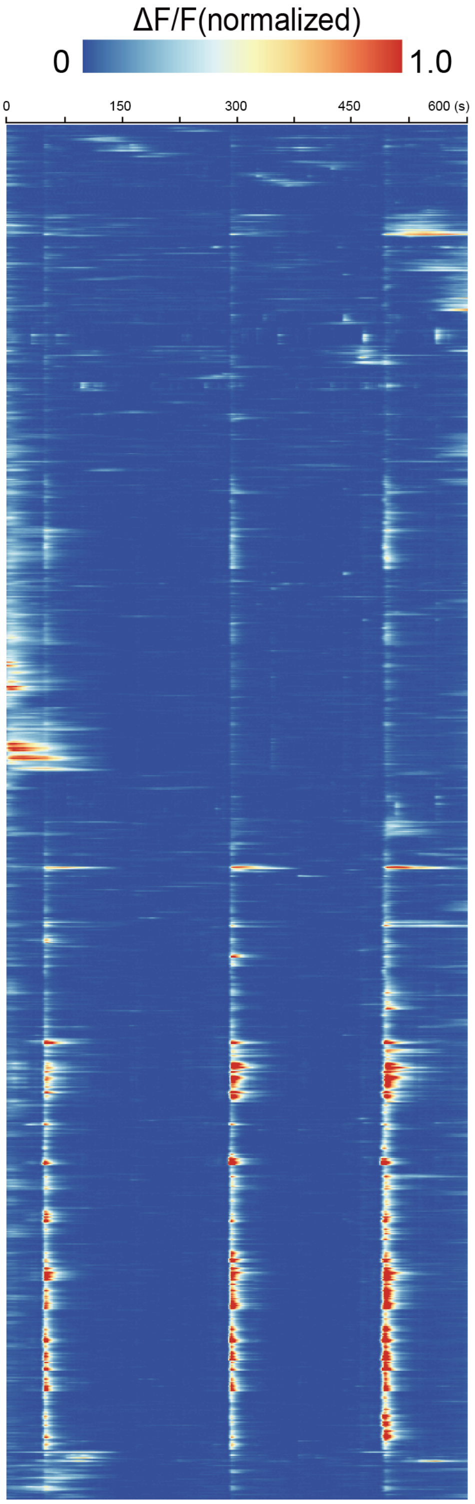


**Supplementary Figure 10. Whole-brain neuronal activity during spontaneous state.**

Heatmap of CLG-calibrated whole-brain neuronal activity during spontaneous behavior; time is shown on the x-axis and each row represents a single neuron.

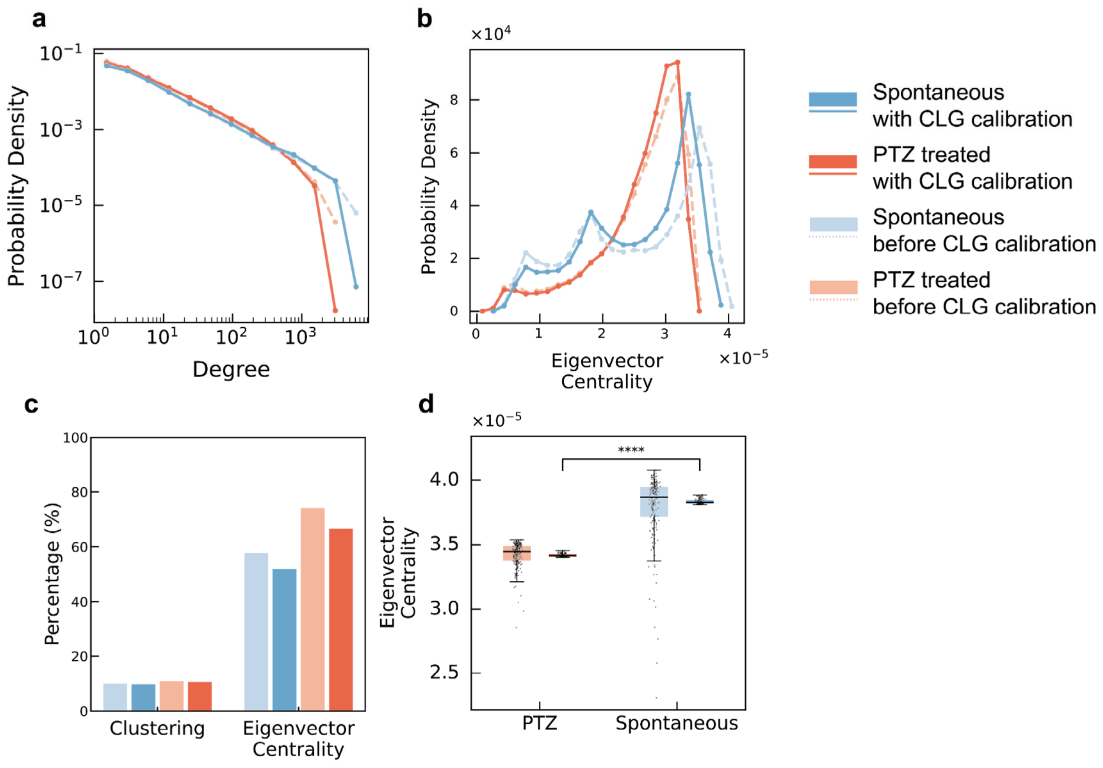


**Supplementary Figure 11. Network analysis before and after CLG calibration.**

**a, b.** Degree and eigenvector centrality distributions for correlation-based networks before (dashed) and after (solid) calibration (KS test, $p < 0.0001$). **c,** Overlap of the top 1,000 nodes ranked by clustering coefficient (left) or eigenvector centrality (right) between pre- and post-calibration networks. **d,** Eigenvector centrality values in the pre-calibration network for the top 100 nodes identified after calibration.

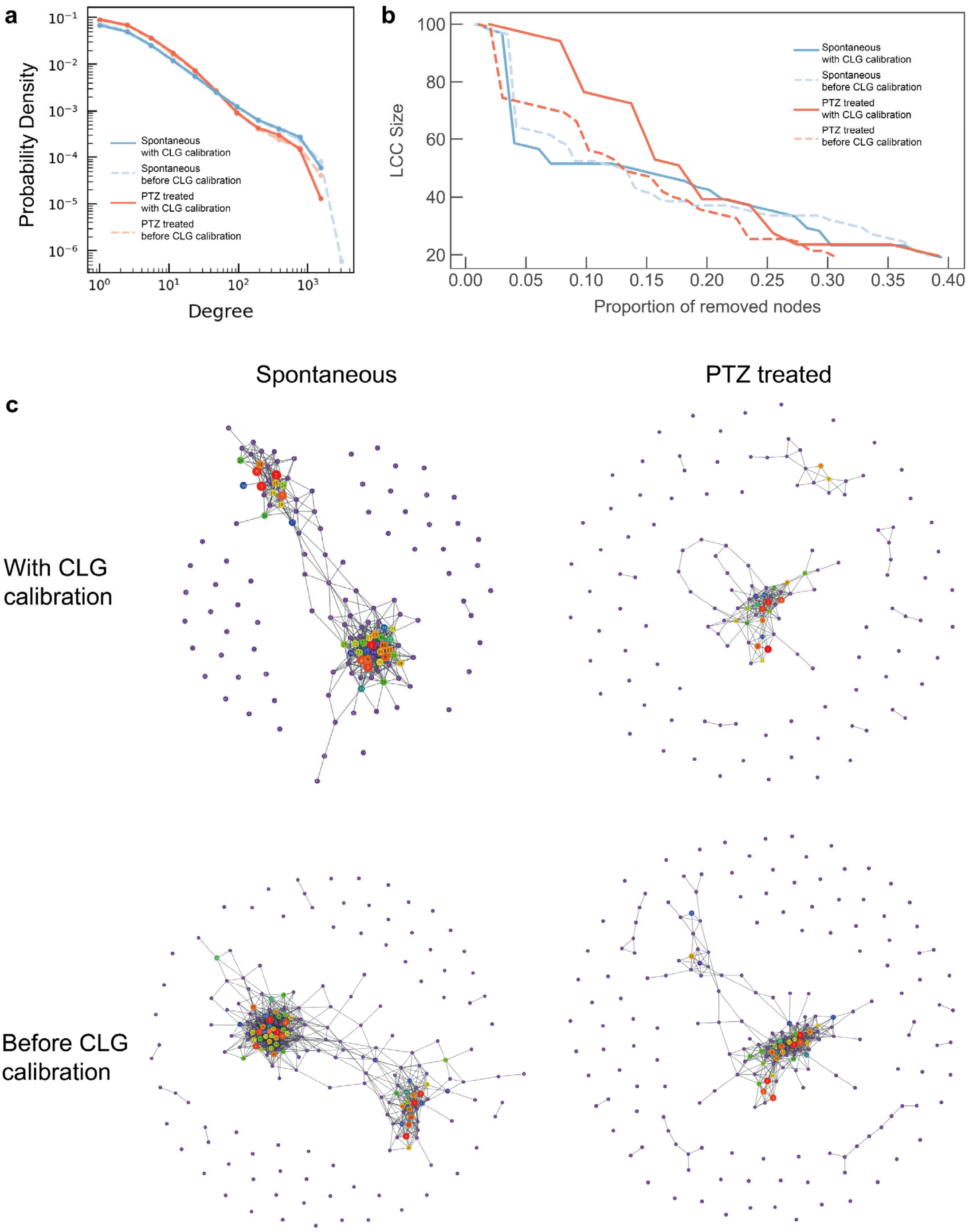


**Supplementary Figure 12. Network topology analysis for Biological Replicate 2 (PTZ-Fish 2).**

**a,** Degree distribution. **b,** Network dismantling analysis using the adapted GDM algorithm. **c,** Visualization of coarse-grained networks for spontaneous and PTZ-treated states before and after calibration.

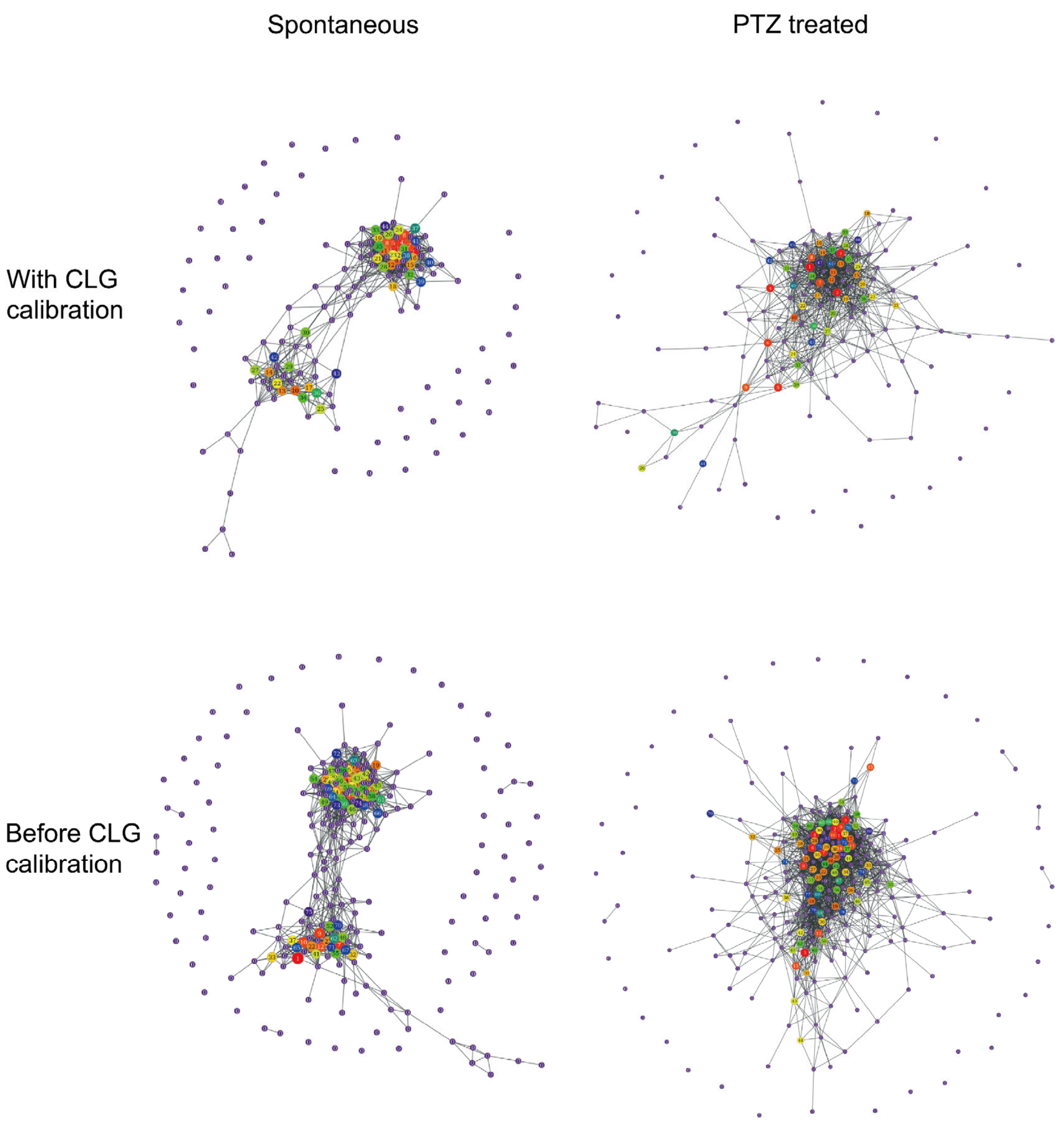


**Supplementary Figure 13. Visualization of coarse-grained networks.**

Direct visualization of coarse-grained networks for spontaneous and PTZ-treated states before and after calibration.

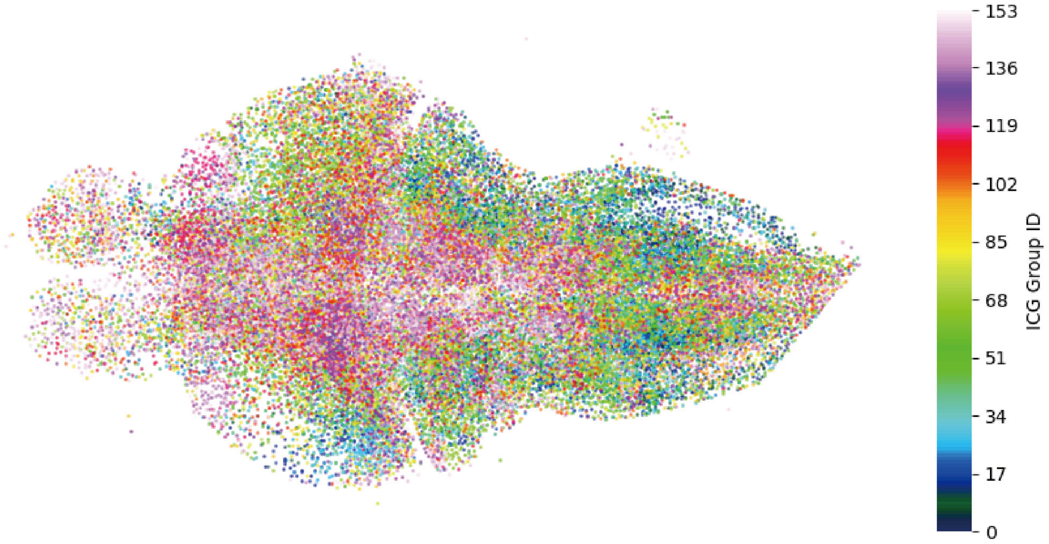


**Supplementary Figure 14. Spatial distribution of coarse-grained network nodes during spontaneous activity.**

Spatial organization of correlation-based coarse-grained nodes (256 cells per node), with each color representing a node.

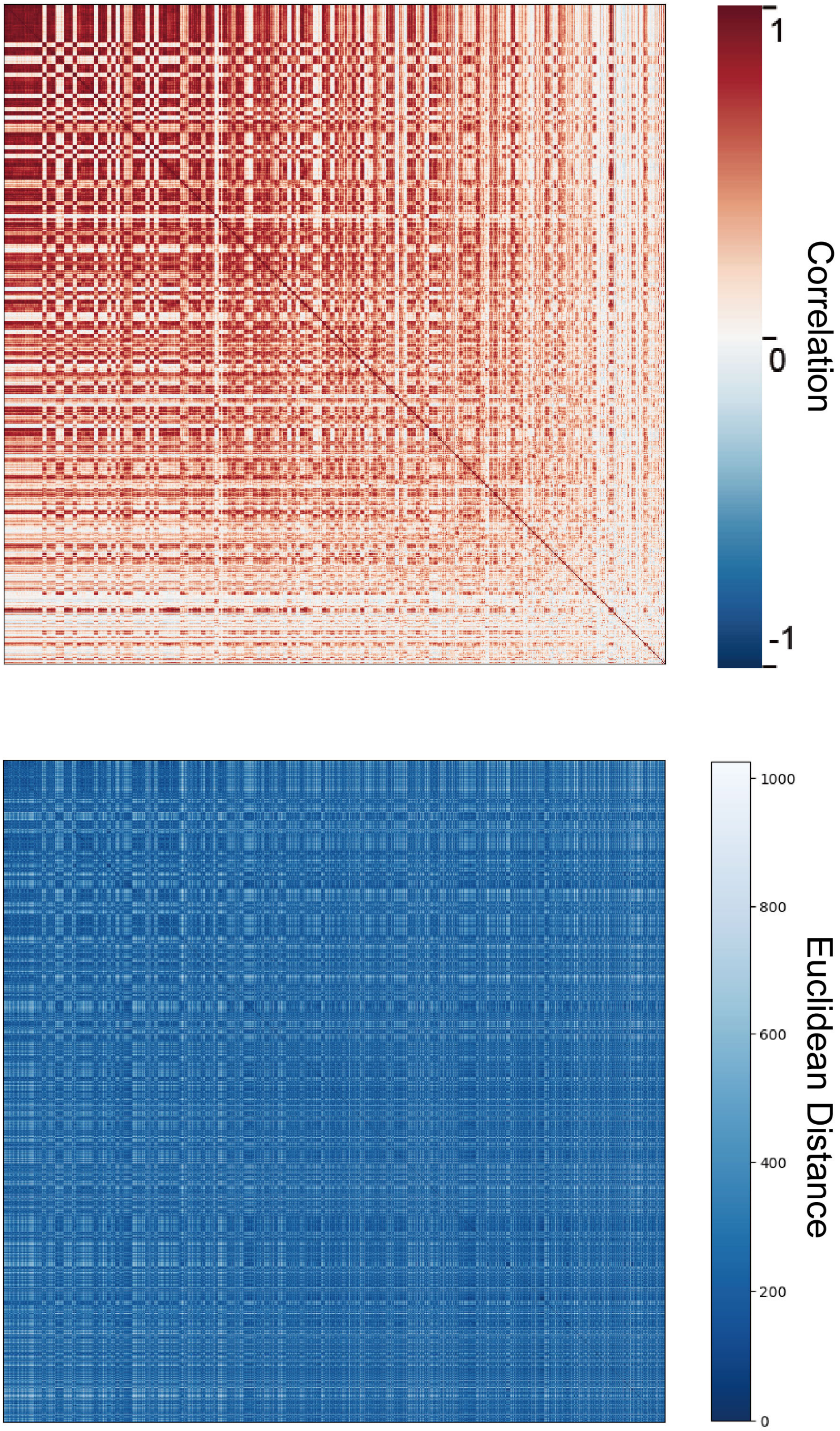


**Supplementary Figure 15. Pairwise correlation and spatial distance matrices during spontaneous activity.**

Pairwise correlation and Euclidean distance matrices ordered by coarse-grained groups.

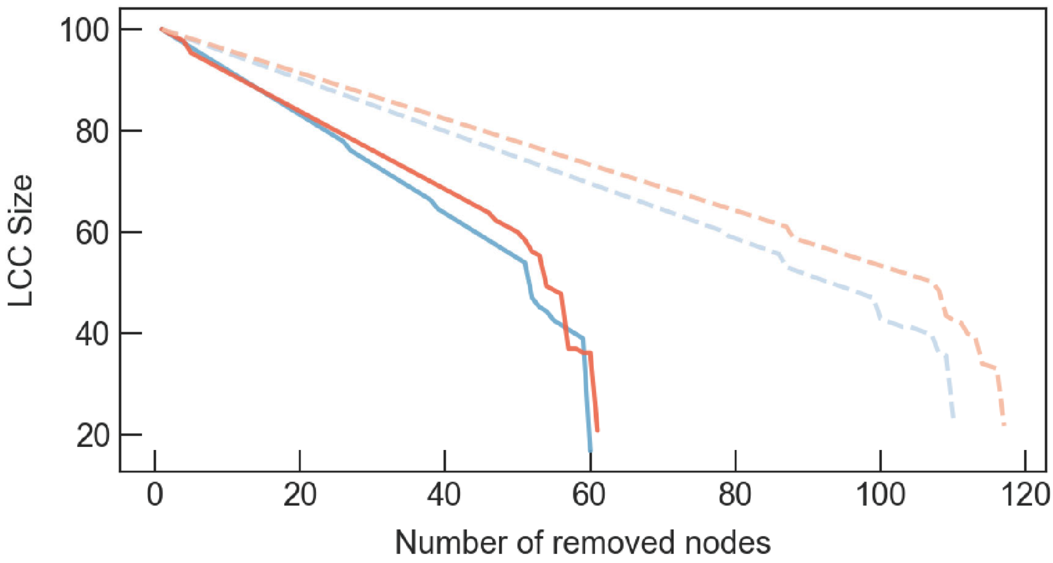


**Supplementary Figure 16. Network dismantling using the CoreHD algorithm.**

LCC size as a function of the number of removed nodes.

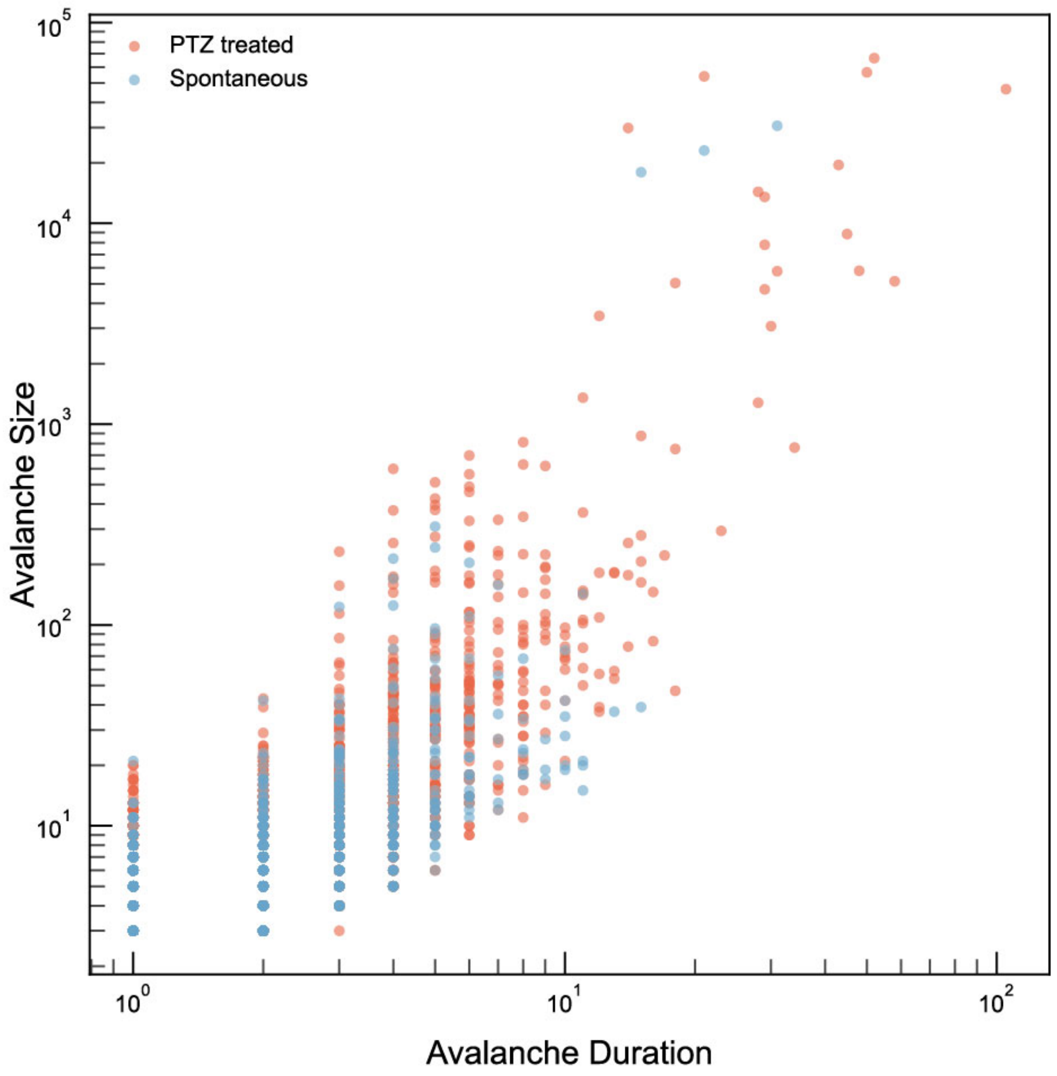


**Supplementary Figure 17. The scaling relationship between avalanche size and duration.**

Each point represents one avalanche event.

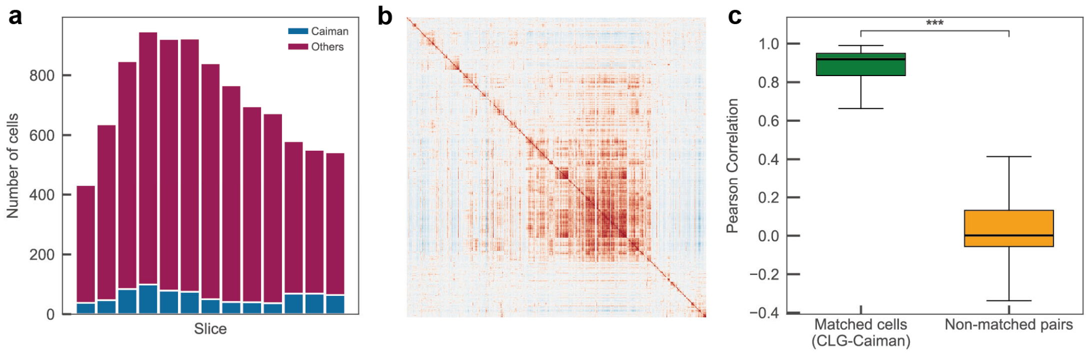


**Supplementary Figure 18. Comparison of CLG- and CaImAn-derived neuron detection and trace fidelity.**

**a,** Neurons identified per imaging slice, partitioned into CaImAn-detected (blue) and CaImAn-missed neurons (maroon). **b,** Pairwise Pearson correlation matrix for activity traces of neurons identified by both CLG and CaImAn. **c,** Quantitative comparison showing significantly higher correlations for matched CLG–CaImAn pairs than unmatched pairs (Mann–Whitney U test, *** $p < 0.001$).

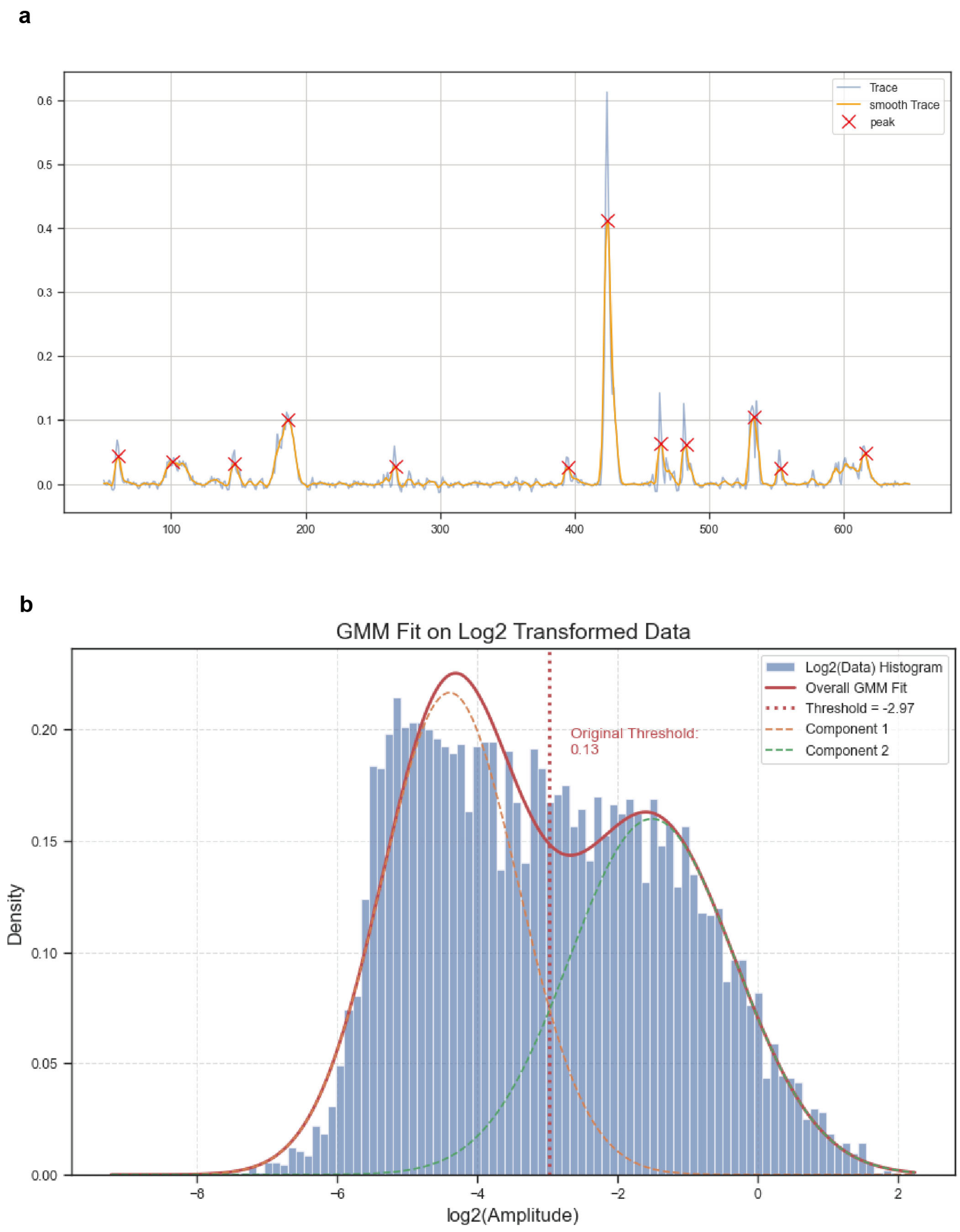


**Supplementary Figure 19. Two-component Gaussian mixture modeling of Δ*F*/*F* distributions.**

**a,** Example of peak detection in Δ*F*/*F* activity. **b,** Two-component Gaussian mixture fit used to define the activity threshold (Δ*F*/*F* > 0.13).

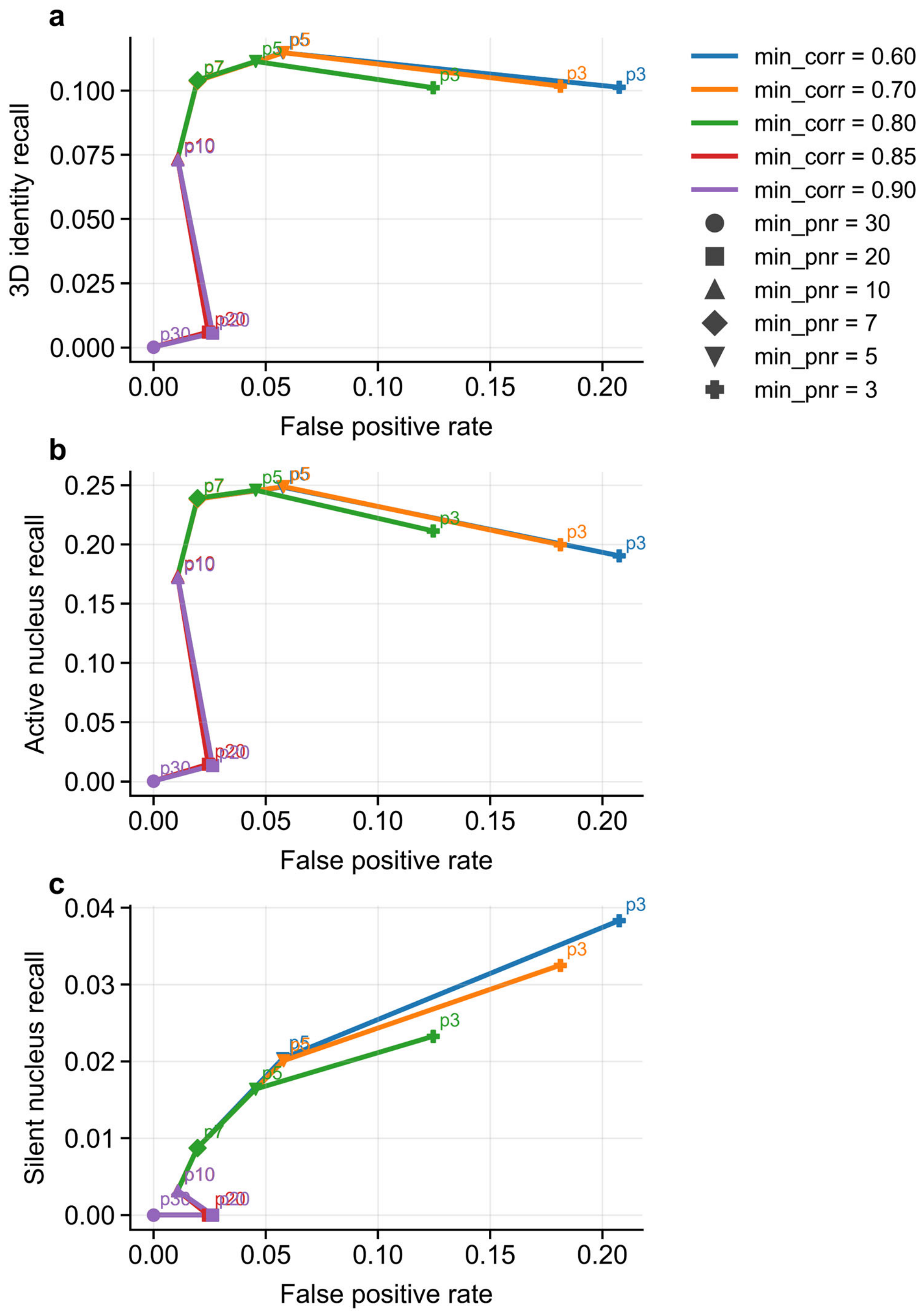


**Supplementary Fig 20. CaImAn parameter titration against nucleus-defined ground truth in mouse visual cortex.**

a–c, CaImAn performance across 20 tested initialization-parameter combinations at fixed gSig=8, plotted as recall versus false positive rate. a, 3D identity recall, defined as the fraction of CLG-defined unique nucleus identities recovered by CaImAn after CLG calibration. b, Active nucleus recall, computed for nuclei classified as active using the same ΔF/F>0.13 threshold as in the main analysis. c, Silent nucleus recall, computed for nuclei below this activity threshold. Curves connect tested settings with the same min_corr, marker shapes indicate min_pnr, and point labels show min_pnr values. Tested combinations were: min_corr=0.60 with min_pnr=3,5,7,10; min_corr=0.70 with min_pnr=3,5,7,10; min_corr=0.80 with min_pnr=3,5,7,10,20,30; min_corr=0.85 with min_pnr=10,20,30; and min_corr=0.90 with min_pnr=10,20,30. False positive rate was defined as the fraction of CaImAn components whose nearest nucleus exceeded the 30-pixel ground-truth matching cutoff. Relaxing min_pnr increased identity-level and active-nucleus recall over an intermediate range, whereas more permissive settings increased false positives without further improving recovery; silent-nucleus recall remained low across settings.

#### 2. Supplementary Video Legends

**Supplementary Video 1. Whole-brain nuclear structural imaging in zebrafish larvae.** Three-dimensional structural imaging of the entire brain in dual-labeled zebrafish larvae expressing H2B-mRuby3.

**Supplementary Video 2. Orthogonal views of whole-brain functional calcium imaging.** Three orthogonal views of GCaMP6s signals illustrating the 3D distribution and dynamics of neuronal calcium activity.

**Supplementary Video 3. Single-plane nuclear imaging and segmentation pipeline.** Workflow from raw nuclear images through preprocessing (sparse deconvolution and local normalization) to final nuclear segmentation.

**Supplementary Video 4. Whole-brain 3D nuclear identification and reconstruction.** Three-dimensional reconstruction of all identified neuronal nuclei, with each nucleus represented as a sphere.

**Supplementary Video 5. Integrated 3D reconstruction of nuclei and single-cell activity.** Combined visualization of nuclear positions and corresponding calcium activity traces across the zebrafish brain.

**Supplementary Video 6. Structural–functional colocalization.** Demonstration of spatial alignment between functional calcium signals (GCaMP6s, green) and nuclear structural signals (mRuby3, red).

**Supplementary Video 7. Cells spanning multiple functional imaging planes.** Three-dimensional reconstruction showing neurons spanning multiple axial planes, corresponding to **Fig. 2b**.

**Supplementary Video 8. Slice-by-slice scanning of whole-brain 3D single-cell identification.** Schematic visualization of sequential scanning and volumetric single-cell identification in spontaneous and PTZ-treated experiments.

**Supplementary Video 9. Whole-brain activity under PTZ-induced epileptic state.** Three-dimensional reconstruction of spatiotemporal neuronal activity 30 min after PTZ administration.

**Supplementary Video 10. Whole-brain activity during spontaneous state.** Three-dimensional reconstruction of baseline neuronal activity across the zebrafish brain.

**Supplementary Video 11. Dual-channel 3D imaging in the mouse cortex.** Three-dimensional reconstruction of mRuby3-labeled nuclei and GCaMP6s cytosolic signals in mouse visual cortex ($510 \times 510 \times 260\ \mu m^3$).

**Supplementary Video 12. Slice-by-slice visualization of 3D single-cell identification in mouse brain.** Sequential visualization of single-cell identification across imaging planes.

**Supplementary Video 13. Single nucleus spanning multiple axial planes in mouse brain.** 3D reconstruction of a representative nucleus extending across multiple planes with 20 μm spacing.

**Supplementary Video 14. 3D nuclear identification in the mouse brain.** Three-dimensional reconstruction of identified neuronal nuclei, represented as spheres.

**Supplementary Video 15. 3D reconstruction of single-cell activity in the mouse brain.** Integrated visualization of nuclear positions and extracted single-cell calcium activity traces.
